# On the Geometry of Somatosensory Representations in the Cortex

**DOI:** 10.1101/2024.07.11.603013

**Authors:** Noam Saadon-Grosman, Tsahi Asher, Yonatan Loewenstein

## Abstract

It is well-known that cortical areas specializing in the processing of somatosensory information from different parts of the body are arranged in an orderly manner along the cortex. It is also generally accepted that in the cortex, somatosensory information is initially processed in the primary somatosensory cortex and from there, it is hierarchically processed in other cortical regions. Previous studies have focused on the organization of representation at a level of a single or few cortical regions, identifying multiple body maps. However, the question of the large-scale organization of these different maps, and their relation to the hierarchical organization has received little attention. This is primarily because the highly convoluted shape of the cortical surface makes it difficult to characterize the relationship between cortical areas that are centimeters apart. Here, we used functional MRI to characterize cortical responses to full-body light touch stimulation. Our results indicate that the organization of both body representation and hierarchy is radial, with a small number of extrema that reign over a large number of cortical regions. Quantitatively computing the local relationship between the gradients of body and hierarchy maps, we show that the interaction between these two radial geometries, body representation and hierarchy in S1 are approximately orthogonal. However, this orthogonality is restricted to S1. Similar organizational patterns in the visual and auditory systems suggest that radial topography may be a common feature across sensory systems.

**Significance statement:** The sensation of touch on our skin is represented in the brain as a map, where body parts are organized sequentially from head to toe. In the cerebral cortex, multiple body maps are distributed across numerous regions, processing signals at different hierarchical levels. Is there a large-scale organization of these body maps in the cerebral cortex? We show that all previously known body maps and their hierarchies are organized with a radial geometry. Similar radial geometry may also characterize the visual and auditory systems, indicating that radial geometry is a common organizational principle of sensory processing in the cortex.

## Introduction

The body map representation in the post-central gyrus of the cerebral cortex is one of the best-known findings in neuroscience. Penfield and Boldrey found that an electrical stimulation in this area, later known as the primary somatosensory cortex (S1, Brodmann areas 3a,3b,1 and 2), elicits a localized somatosensory sensation (Penfield and Boldrey 1937). Moving the stimulating electrode from the lateral to the medial part of the gyrus, resulted in sensation moving from the head to the toes, along the long axis of the body (Penfield and Boldrey 1937; Penfield and Jasper 1954; Penfield and Rasmussen 1950). In line with these findings, later studies showed that somatosensory stimulations of different parts of the body in both humans and non-human primates elicit neural activity in the corresponding S1 subregions (Fox et al. 1987; Kaas et al. 1979; Merzenich et al. 1978; Muret et al. 2022; Nakamura et al. 1998; Saadon-Grosman et al. 2020b; Sanchez Panchuelo et al. 2018; Tal et al. 2017).

Going beyond S1, Penfield and his colleagues also identified additional body-map representations in cortical regions outside S1 (Penfield 1950; Penfield and Boldrey 1937; Penfield and Faulk 1955; Penfield and Jasper 1954). Later studies confirmed many of Penfield’s findings and identified even more somatosensory body-map representations (Arienzo et al. 2006; Burton et al. 1993; Fitzgerald et al. 2004; Fox et al. 1987; Huang et al. 2012; Kaas et al. 1979; Kaas and Collins 2001; Lim et al. 1994; Nakamura et al. 1998; Ruben et al. 2001; Saadon-Grosman et al. 2020b; Sakata et al. 1973; Sanchez Panchuelo et al. 2018; Young et al. 2004).

These body maps describe the functional organization of the cortex within each of these cortical regions. However, it has not been clear what principles underlie the functional organization of the cortex at the larger scale of multiple cortical regions.

For comparison, in the visual modality, the visual field is retinotopically mapped within cortical regions in two dimensions-polar angle, and eccentricity. On a scale larger than a single cortical region, these individual maps are organized into several clusters, groups of maps with parallel semicircular polar angle representations that share a common eccentricity representation (Wandell et al. 2005, 2007). The large-scale organization in the visual domain has motivated us to study the macroscopic organization of maps in the somatosensory cortex.

The body map representation alone does not capture all characteristics of neural responses. In S1, electrophysiological recordings in non-human primates have revealed that as we move away from the central sulcus caudally, the receptive fields of the neurons increase in size and complexity (Hyvärinen and Poranen 1978; Iwamura 1998; Iwamura et al. 1993; Taoka et al. 1998). Similarly, BOLD responses in area 3b were found to be more selective than those in area 1, which in turn, were more selective than those in area 2 (Ann Stringer et al. 2014; Puckett et al. 2020; Saadon-Grosman et al. 2020a; Schellekens et al. 2021). The increase in size and complexity of the receptive fields are indicative of a rostral to caudal hierarchical processing direction in S1. This hypothesis is supported by the finding that neural response latencies increase in that direction (Inui et al. 2004; Lebedev and Nelson 1996) and by anatomical evidence of serial corticocortical connections (Felleman and Van Essen 1991; Vogt and Pandya 1978) and thalamic projections (Jones and Powell 1970). Another measure of the somatosensory hierarchy is the extent to which the neural response is contralateral. As expected, both electrophysiological recordings and magnetic imaging reveal that neural responses become more bilateral as we move in the rostral-caudal direction (Iwamura 1998, 2000; Killackey et al. 1983; Saadon-Grosman et al. 2020a).

Hierarchical organization of the somatosensory system, which manifests as increase in latencies, receptive field size, complexity and laterality of somatosensory responses, has also been observed outside S1 (Felleman and Van Essen 1991; Inui et al. 2004; Iwamura 2003; Kaas 1993; Mazzola et al. 2006; Sakata et al. 1973; Schellekens et al. 2018). In a previous study, we identified somatosensory hierarchical gradients from the central sulcus posteriorly, anteriorly and medially (Saadon-Grosman et al. 2020a). Nevertheless, the general organization of hierarchical processing in the somatosensory cortex is still poorly understood.

In this paper we used BOLD responses to full-body somatosensory stimulation to study the large-scale functional organization of the somatosensory cortex. We show that at the largest scale, radial geometry describes well both the organization of body maps and the organization of hierarchical processing. We further show that as the result of the interaction of these two radial geometries, the long axis of the body representation and hierarchy in S1 are approximately orthogonal, allowing S1 to process somatosensory information from different body parts at all levels of hierarchy. However, this orthogonality is not observed outside S1.

## Results

To characterize somatosensory responses, we used fMRI to measure BOLD responses to tactile stimulation along the long axis of the body (Saadon-Grosman et al. 2020a, 2020b) (see Methods). We used these BOLD responses to construct a body map, which depicts the preferred body part at each vertex (Fig. 1A, for comparable anatomy, see Fig. S1A). For clarity, we present, throughout the manuscript, only the left hemisphere. Comparable figures for the right hemisphere are presented in the Supporting information section (Figs. S2A and S3A).

**Figure 1.**
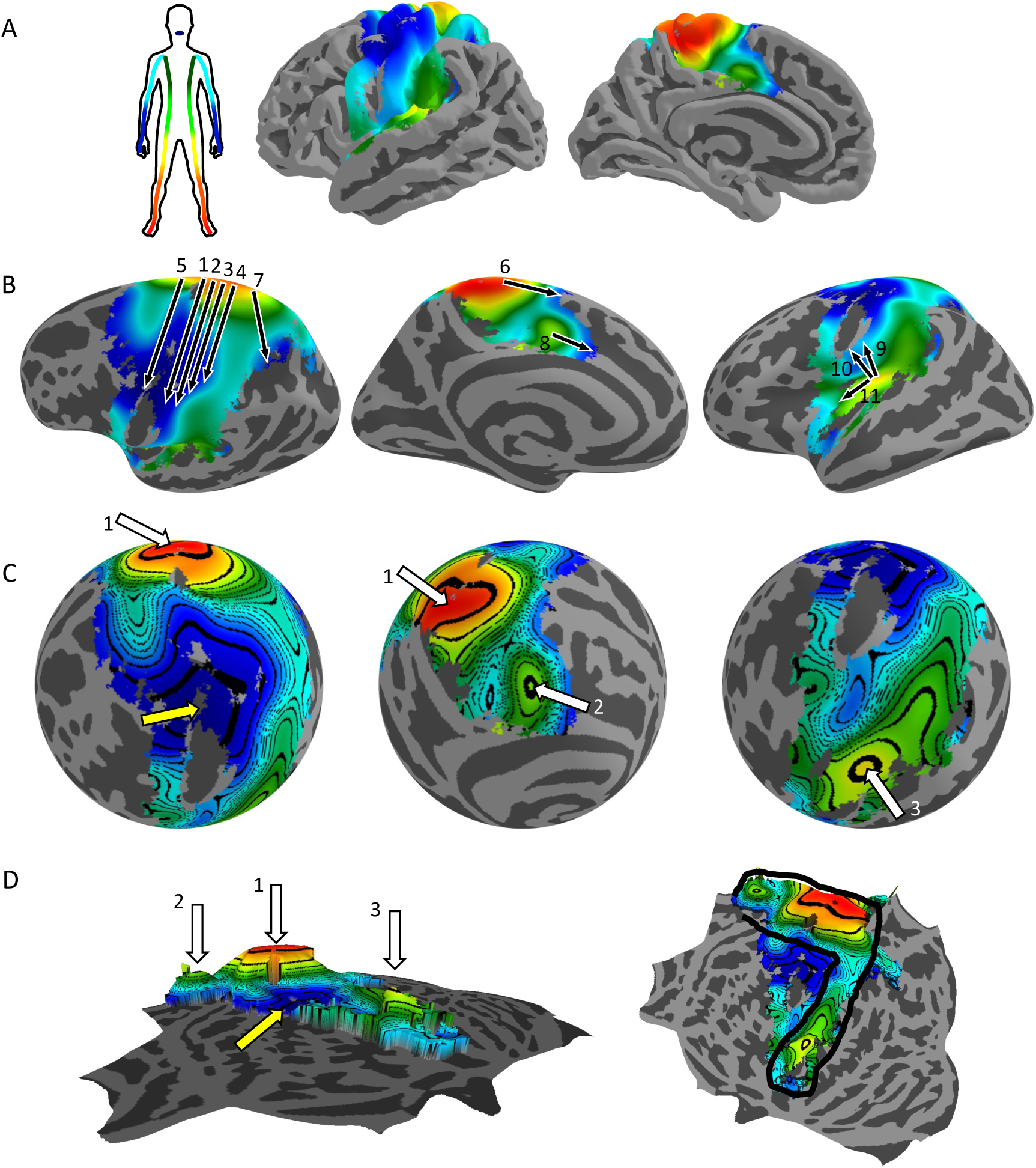
Radial topography of somatosensory body representation. A group (N=20) body map corresponding to the stimulation of the contralateral body side, from lips (blue) to toes (red), is displayed on multiple representations of the left hemisphere cortical surface. A. Pial surface in lateral (left) and medial (right) views. B. Inflated surface in lateral (left), medial (middle), and inferolateral (right) views. Black arrows mark known body maps from previous literature (see text), note their consistency with the underlying map. C. Spherical representation of the cortical surface in three views. Contour lines represent vertices corresponding to the same body part. D. Flat surface representation, showing a side view where the body part value determines the height above the surface (left), and a top view of the same representation (right). The body map is characterized by a single minimum in the lateral part of the pre- and post-central gyri (lips, yellow arrow in C and D), surrounded by a semicircular ridge (black line in D) that spans the posterior extent of the entire somatosensory cortex on the medial and lateral surfaces. The ridge consists of three maxima (leg representation). The largest of these maxima lies at the midline, between the medial and lateral surfaces of the hemisphere, within the medial extent of the pre- and post-central gyri (arrow 1). A second, elongated peak is located deep within the Sylvian fissure/posterior insula (arrow 3). The third maximum is found in the cingulate gyrus on the opposite side of the ridge (arrow 2).

Body map representations have been studied from the time of Penfield and onwards using different techniques. Specifically, 11 complete maps have been reported, and we start by showing that they are consistent with our measurements. Fig. 1B depicts these maps (arrows) on an inflated cortical representation (Fig. 1B, Figs. S1-3B). There are four body maps, which align with the four anatomical areas in the postcentral gyrus (Brodmann 3a, 3b, 1, and 2)(Blankenburg et al. 2003; Gelnar et al. 1998; Kaas et al. 1979; Sánchez-Panchuelo et al. 2014). These are depicted by arrows 1-4. In these maps, the long axis of the body spans from medial to lateral, ranging from the toes to the lips. Adjacent to S1, anteriorly on the opposite side of the central gyrus, a complete somatosensory body map was found in the primary motor cortex (M1, arrow 5) (Penfield and Boldrey 1937). There are two additional maps that ‘share’ a leg representation with those five maps and ‘meet’ in the medial extent of the pre- and post-central gyri, on the midline between the lateral and medial surfaces of the hemisphere. One map is located within the supplementary motor area (SMA), starting from the hemisphere midline and extending inferiorly and anteriorly on the medial wall (Arienzo et al. 2006; Penfield and Jasper 1954)(arrow 6). The other map is located in the superior posterior parietal cortex (Huang et al. 2012) (arrow 7).

In the medial wall lies another map, which starts above the cingulate cortex and progresses anteriorly (Arienzo et al. 2006) (arrow 8). In the secondary somatosensory cortex in the upper bank of the Sylvian fissure, three body maps have been reported. One starts deep within the Sylvian fissure, spanning superiorly and anteriorly in the parietal opercula (OP1, analogous to S2 in monkeys), and terminating in the lateral extent of S1 (arrow 9). The other, anterior to it, in a similar orientation is in the parietal ventral area (OP4, analogous to PV in monkeys) (arrow 10)(Disbrow et al. 2000; Eickhoff et al. 2007; Krubitzer et al. 1995) (Burton et al. 1993; Fox et al. 1987; Penfield and Jasper 1954). Finally, a body representation that spans from the depth of the Sylvian fissure anteriorly (OP2-3, analogous to VS in monkeys) has been reported (Eickhoff et al. 2007; Penfield and Faulk 1955)(arrow 11). Notably, these previous maps were constructed using different techniques and stimuli in different labs. Our findings, using a single experimental paradigm and analysis, are consistent with all these previous studies.

To better understand the macroscopic organization of these maps, we consider the even more inflated spherical representation of the cortex, in which we also added contour lines that correspond to areas of equal-body representation, similar to a topographic map (Fig. 1C, Figs. S1-3C). Considering this ‘topographic map’, in which the lips correspond to the lowest altitude and the toes to the highest, we see that the body map is characterized by a large minimum in the lateral part of the pre- and post-central gyri (yellow arrow). This minimum is the face-region in many of the discrete maps discussed above, namely arrows 1-5, 9 and 10. Superiorly, inferiorly and posteriorly from this minimum, we observe three maxima (leg representation), which are denoted by white arrows in the figures. The largest of these lies in the midline, between the medial and lateral surfaces of the hemisphere, in the medial extent of the pre- and post-central gyri (white arrow 1). This peak corresponds to the leg representation of maps 1-7 described above. A second, peak lies above the cingulate gyrus, the leg representation of arrow 8 (white arrow 2). Finally, we see a third peak in the depth of the Sylvian fissure / posterior insula. This peak potentially connects the leg representations of arrows 9-11 (white arrow 3).

A more careful analysis of each these peaks reveals that they are elongated. The largest peak (white arrow 1) is elongated in the anterior-posterior direction, the second peak (white arrow 2) is slightly elongated in the same direction whereas the third one is strongly elongated in the superior-inferior direction. To gain better insight to the organization of these peaks, we consider a flat cortical representation (Fig. 1D, Figs. S1-3D), which enables us to view the entire somatosensory cortex on the same plane. Using this representation, it becomes evident that from a bird’s eye view, the three peaks are connected along a “ridge” that spans the posterior extent of the entire somatosensory cortex in the medial and lateral surfaces and forms a semicircular enclosure around a basin. Thus, from the lateral part of the pre- and post-central gyri, which represents the lips, representations smoothly and monotonically change radially along the long axis of the body in all directions in which we find somatosensory representation, namely superiorly, inferiorly and posteriorly. A single full face-to-legs body representation is evident when moving from the lips’ basin in the lateral part of the pre- and post-central gyri to the three peaks of the ridge. Moving from the basin to the saddles yields a partial body representation that contains only the upper parts. The ridge is associated with a representation of the lower parts of the body, where the peaks represent legs reversal axes (mirror reversal) and the saddles represent reversals at intermediate parts of the body. Next, we went beyond the body map representation to consider the hierarchical structure of somatosensory processing. To that goal, we used the fact that while cortical vertices respond maximally to the somatosensory stimulation of one of the body parts, as depicted in Fig. 1 and Fig. S2, they typically also respond, to a lesser extent to the stimulation of adjacent body parts. This allows us to construct a “tuning curve” for each vertex (Methods), and the width of this tuning curve is a measure of the selectivity of the vertex: the larger the width, the less selective is the vertex. In the visual domain, neural responses become less selective as we move from the primary visual cortex to higher cortical regions. Similarly, in artificial convolutional neural networks, neurons become less selective as we move away from the input layer to higher levels (O’Shea and Nash 2015). Therefore, we used the width of the somatosensory tuning curve as a proxy for the processing level. Fig. 2A (Fig. S4A) depicts the selectivity (the reciprocal of the width of the tuning curve) across the cortex. To relate this selectivity to previous studies, it is useful to consider the inflated cortical surface Fig. 2B (Fig. S4B). Most previous studies on hierarchy in somatosensory processing either focused on specific somatosensory areas and characterized hierarchy within them, or measured hierarchy between very specific cortical regions.

**Figure 2.**
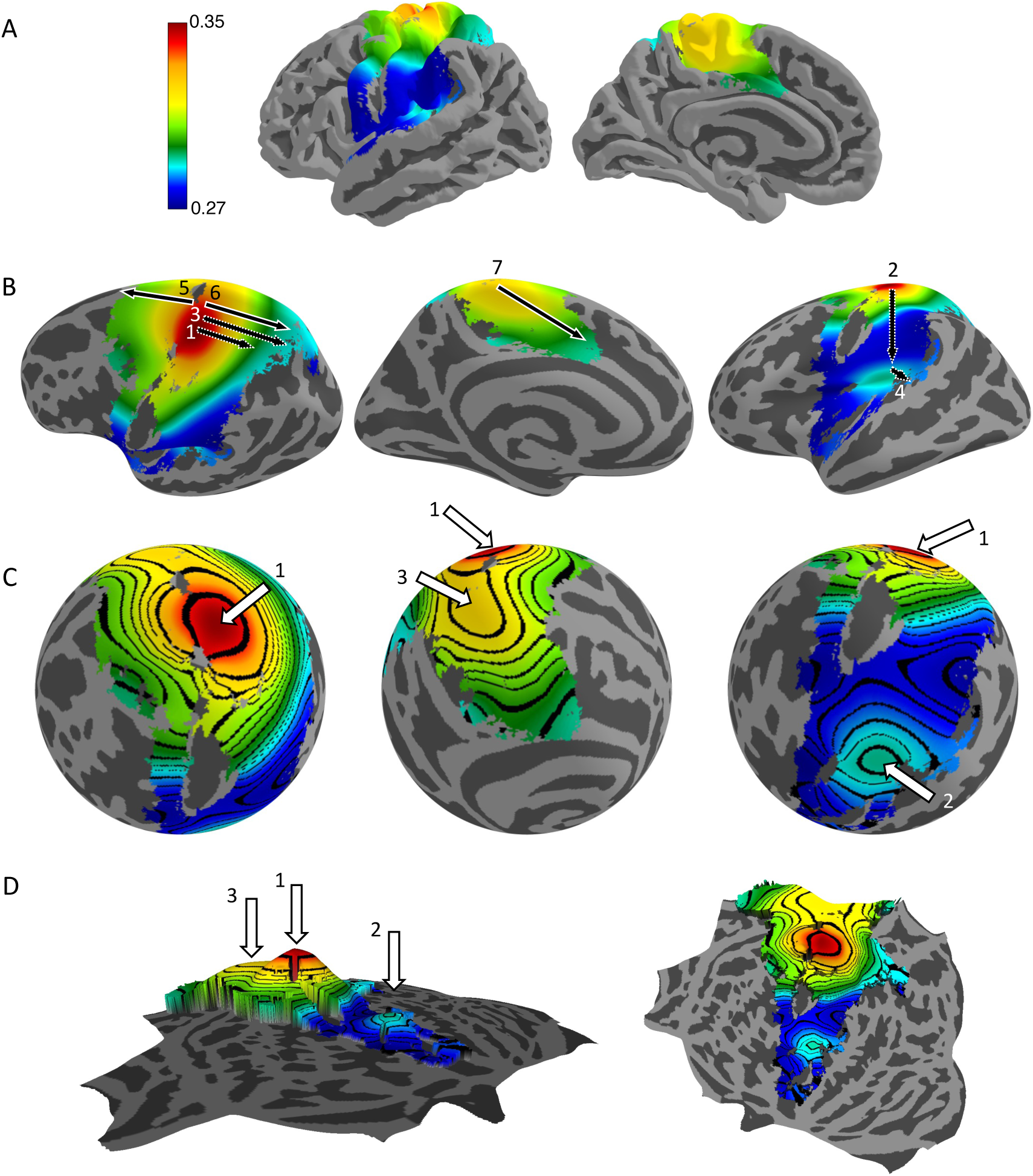
Radial topography of somatosensory processing hierarchy. A group (N=20) selectivity map showing the specificity of the cortical response in each vertex to the preferred body part (high selectivity -red; low selectivity -blue) is displayed on various representations of the left hemisphere cortical surface (same as in Fig. 1). A. Pial surface in lateral (left) and medial (right) views. B. Inflated surface in lateral (left), medial (middle), and inferolateral (right) views. Black arrows indicate known hierarchies from previous literature (see text). Note the consistency of the arrows with the underlying selectivity map. C. Spherical representation of the cortical surface in three views. Contour lines represent vertices of equal selectivity. D. Flat surface representation, showing a side view where the selectivity value determines the height above the surface (left), and a top view of the same representation (right). The selectivity map is dominated by a maximum (a peak) in the anterior part of the postcentral gyrus (arrow 1). There are two additional local maxima of hierarchy: one in the depth of the Sylvian fissure/posterior insula (arrow 2), and one in the superior medial wall (arrow 3).

Within S1, numerous studies have shown that area 3b is the first and main target of afferent thalamic projections of tactile information (Felleman and Van Essen 1991; Iwamura 1998; Iwamura et al. 1993; Killackey et al. 1983). Posterior to it, somatosensory responses in area 1 and then in area 2 exhibit longer latencies, less selectivity and more complexity, all indicating that they are higher the processing hierarchy (Hyvärinen and Poranen 1978; Iwamura 1998, 2003; Iwamura et al. 1993) (arrow 1). Considering the secondary somatosensory area (S2), latencies of responses in this area are longer than in S1 and they are less selective and more bilateral than in S1, indicating that S2 is higher in the processing hierarchy than S1 (Iwamura 2003; Mazzola et al. 2006; Young et al. 2004) (arrow 2). Similar evidence indicates that the posterior parietal cortex is also higher in the processing hierarchy than S1 (arrow 3) (Duffy and Burchfiel 1971; Sakata et al. 1973). Additionally, a previous study has reported hierarchy from S2 to the posterior insular cortex (Mazzola et al. 2006) (arrow 4). Finally, in our previous work, we suggested three streams of descending hierarchy from S1 anteriorly to the frontal lobe (arrow 5), posteriorly to the parietal lobe (arrow 6) and inferiorly in the medial wall (arrow 7) (Saadon-Grosman et al. 2020a).

As with the body map, the largest-scale organization of the map can be better understood when considering the spherical (Fig. 2C, Fig. S4C) and flat (Fig. 2D, Fig. S4D) surface visualizations. With these, it becomes apparent that the selectivity map exhibits a radial geometry with three main peaks (white arrows): (1) a peak in the rostral part of S1 (the border between area 3a and area 3b), more than half way on the lateral-medial axis, (2) a peak in the depth of the Sylvian fissure / posterior insula, and (3) a smaller peak in the superior part of the medial wall, in line with the central sulcus.

So far, we used the width of the somatosensory tuning curve as a proxy for the processing level. There is a different measure of selectivity, which is also indicative of the hierarchy processing level. The afferent pathway to area 3b conveys tactile information that is primarily contralateral, which manifest in contralateral cortical responses. Cortical responses in other areas are less lateralized, and this laterality can be used as a different measure of hierarchy (see Methods). The laterality cortical maps, depicted in Figs. S5 and S6, are remarkably similar to the selectivity maps. Note that while the selectivity in a vertex measures the relative responses of that vertex to somatosensory stimulations of the contralateral body side, the laterality measures the relative responses of that vertex to stimulations of the contralateral and ipsilateral body sides. Because ipsilateral responses were not utilized for constructing the selectivity map, the similarity between the selectivity and laterality maps is an additional indication that selectivity is a useful measure of the level of hierarchical processing.

Having characterized the large-scale organization of body and hierarchy maps, we set out to study how the two maps relate. Specifically, we considered the relationship between the gradients of the body and selectivity maps. The gradient of a map is the direction of the greatest rate of increase along the cortical surface and is orthogonal to the contour lines. The angle between the gradients of the body and selectivity maps is an interesting quantity because it quantified the extent to which the two are locally, independent quantities. If the two gradients are orthogonal then locally, information about the same body part is processed at different level of selectivity, which implies that they are processed at different levels of hierarchy. Conversely, if the gradients are parallel then the hierarchy level is a property of the body part.

We focus on S1, the primary cortical region for somatosensory information processing. The body map in S1 is dominated by the highest peak of the map, leg representation in the medial extent of the pre- and post-central gyri and by the basin, the lips representation in the lateral part of the pre- and post-central gyri (Fig. 3, top, left, Fig.S7). Continuity of body parts’ representation dictates that any line connecting the two peaks should contain all body parts’ representations. To use an analogy from electricity, we can parallel the maximum and the minimum of the body map to positive and negative charges that induce an electric field, such that any line between the two charges contain all the voltages between them. In electricity, a finite dipole consists of two, spatially separated opposite charges. The gradient of the body map is thus similar to the electric field of the dipole, with a pattern of field lines extending outwards from the positive charge to the negative charge (Fig. 3, bottom, left).

**Figure 3.**
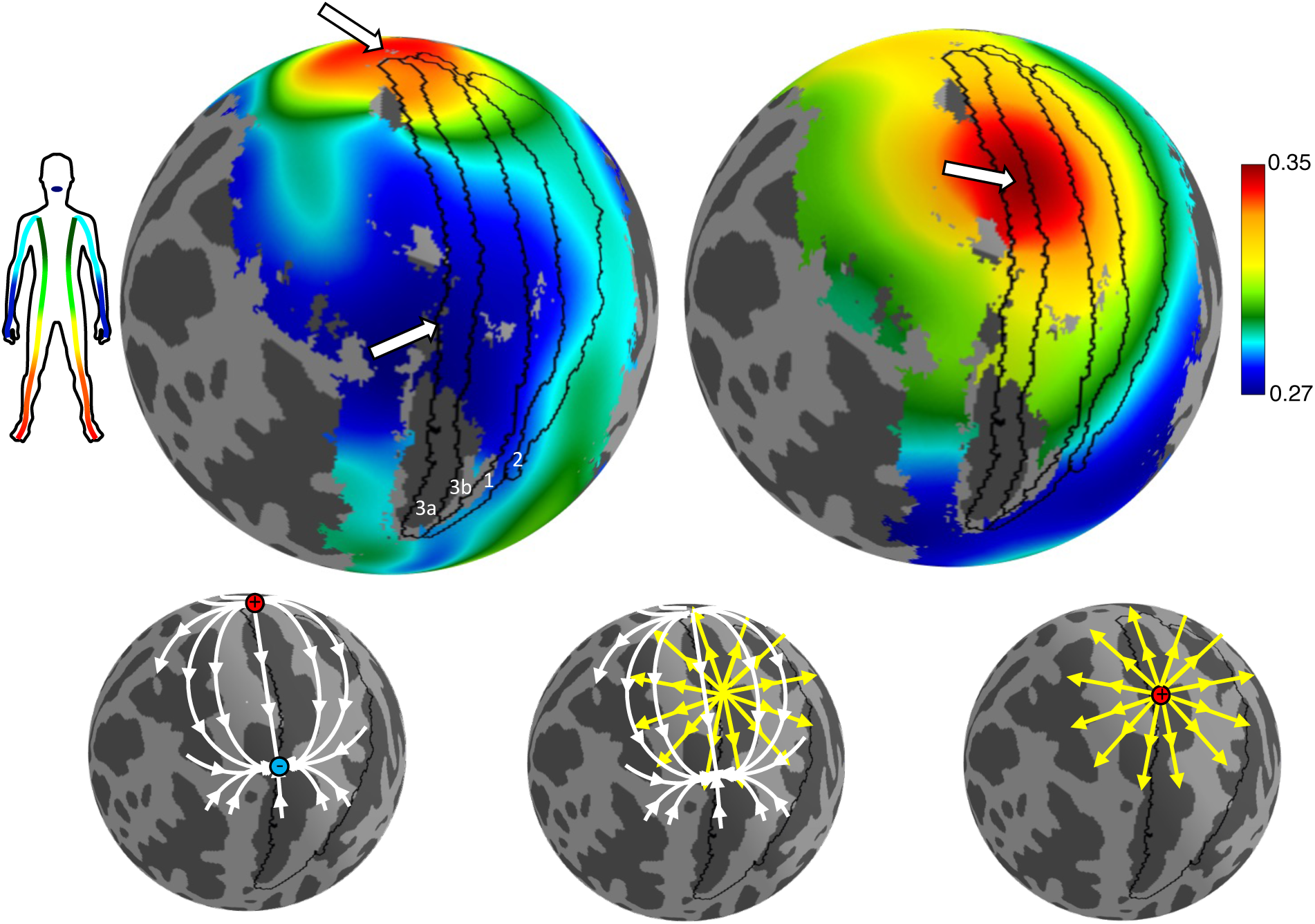
Body representation and hierarchy in the primary somatosensory Cortex (S1) Body map (top left) and selectivity map (top right) are displayed on a spherical representation of the left hemisphere cortical surface in a superior lateral view. Black lines denote the borders of Brodmann areas that comprise S1 (3a, 3b, 1, 2). Thick white arrows mark extrema. In the body map, a maximum (leg representation) is situated near the medial extent of S1, and a minimum (lips representation) is found in the lateral part of S1. In the selectivity map, a maximum is located approximately halfway between these body extrema. To address the geometrical relationship between the gradients of both maps, we use an analogy from electricity. The body extrema can be described as an electric dipole (bottom left) and the selectivity extrema as a point charge (bottom right). The relationship between the body and selectivity gradients resembles those between the electric fields induced by a dipole and by an electric charge (bottom middle).

The selectivity map in S1 is dominated by one peak, in the rostral part of S1, approximately half way between the maximum and minimum of the body maps (Fig. 3, top, right, Fig. S7). Continuing with the electric field analogy, the selectivity map can be paralleled to a point charge, from which the electric field lines radiate symmetrically in all directions (Fig. 3, bottom, right). Therefore, in S1, the relationship between the body and selectivity gradients resembles those between the electric fields induced by a dipole, and those induced by an electric charge (Fig. 3, bottom, middle). Area S1, encompasses almost all of the dipole, and regions posterior and lateral to it. The electric fields analogy predicts that throughout most of S1, the gradient lines of the body and selectivity maps would be approximately orthogonal.

This analysis, depicted in Fig. 3, is based on the spherical visualization of the cortex, which substantially distorts the anatomical relation between cortical areas. Therefore, to quantitatively compute the local relationship between the gradients of the two maps, we developed a method that transforms any scalar map into a vector field on the cortical surface, taking into account the highly non-Euclidian shape of the cortex. In short, at each vertex, we identify the tangent plane to that vertex (Fig. 4A, B).

**Figure 4.**
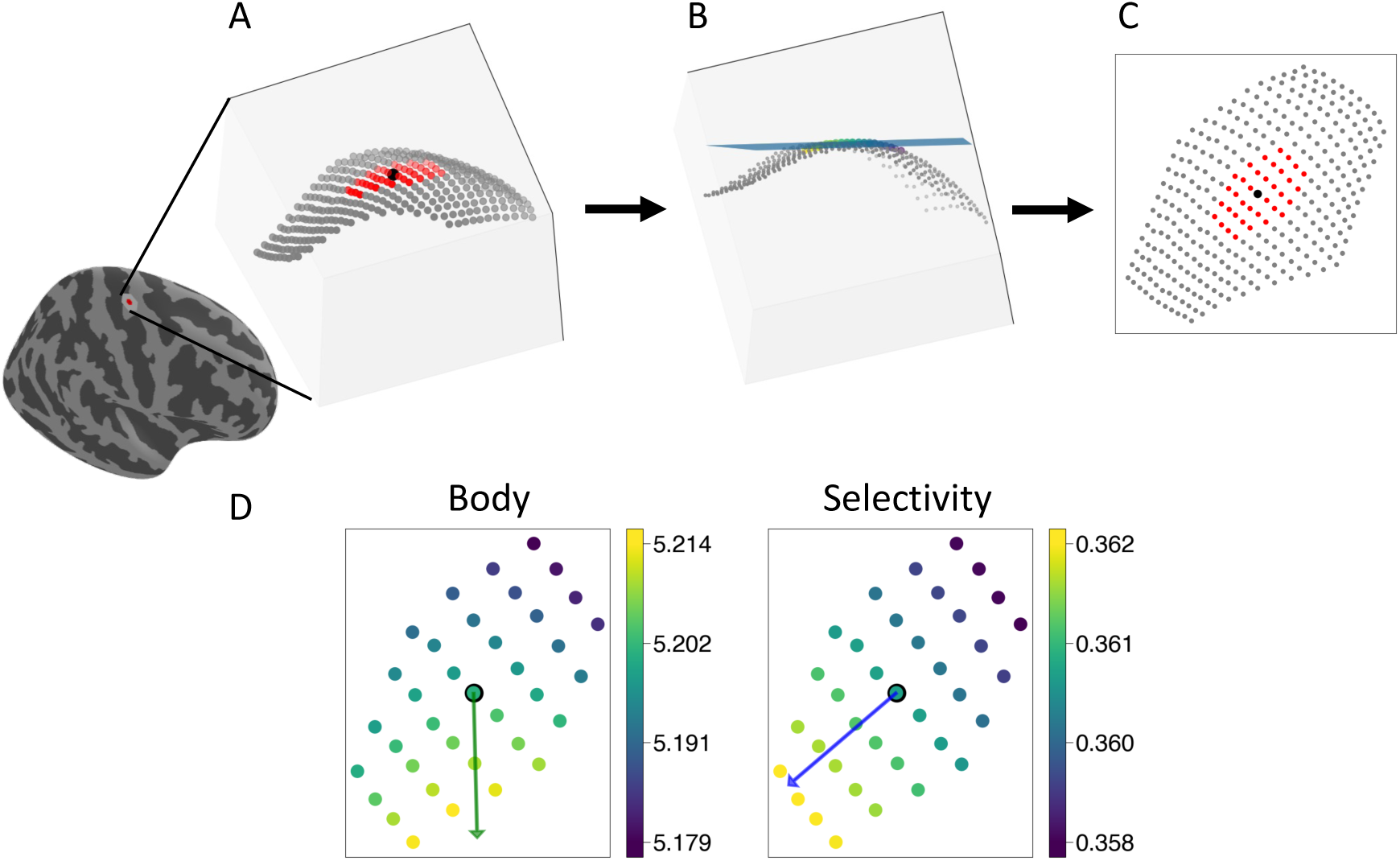
Computing vector fields (gradients) A. To compute the gradient of the map at a vertex, we consider all its third-degree neighboring vertices (red dots). B. First, we use its nearest neighbors in order to identify the plane that is tangent to that vertex. C. Then, we project the vertices on that plane. D. This results in two local maps for body and selectivity (color coded). We then use linear regression to identify the local gradient for each of these maps. The best-fitting line is plotted for each map.

The neighboring vertices are then projected onto that plane (Fig. 4C), allowing us to compute a local gradient of that map (Fig. 4D). This analysis results with, for each vertex, a gradient direction for the body and selectivity maps (Fig. 4D). To test our method, we verified that the gradients of our two measures of hierarchy, selectivity and laterality, are indeed parallel (Figs. S8 and S9). Next, we measured the angles between the gradients of the body and selectivity maps (Figs. 5A, S10A).

**Figure 5.**
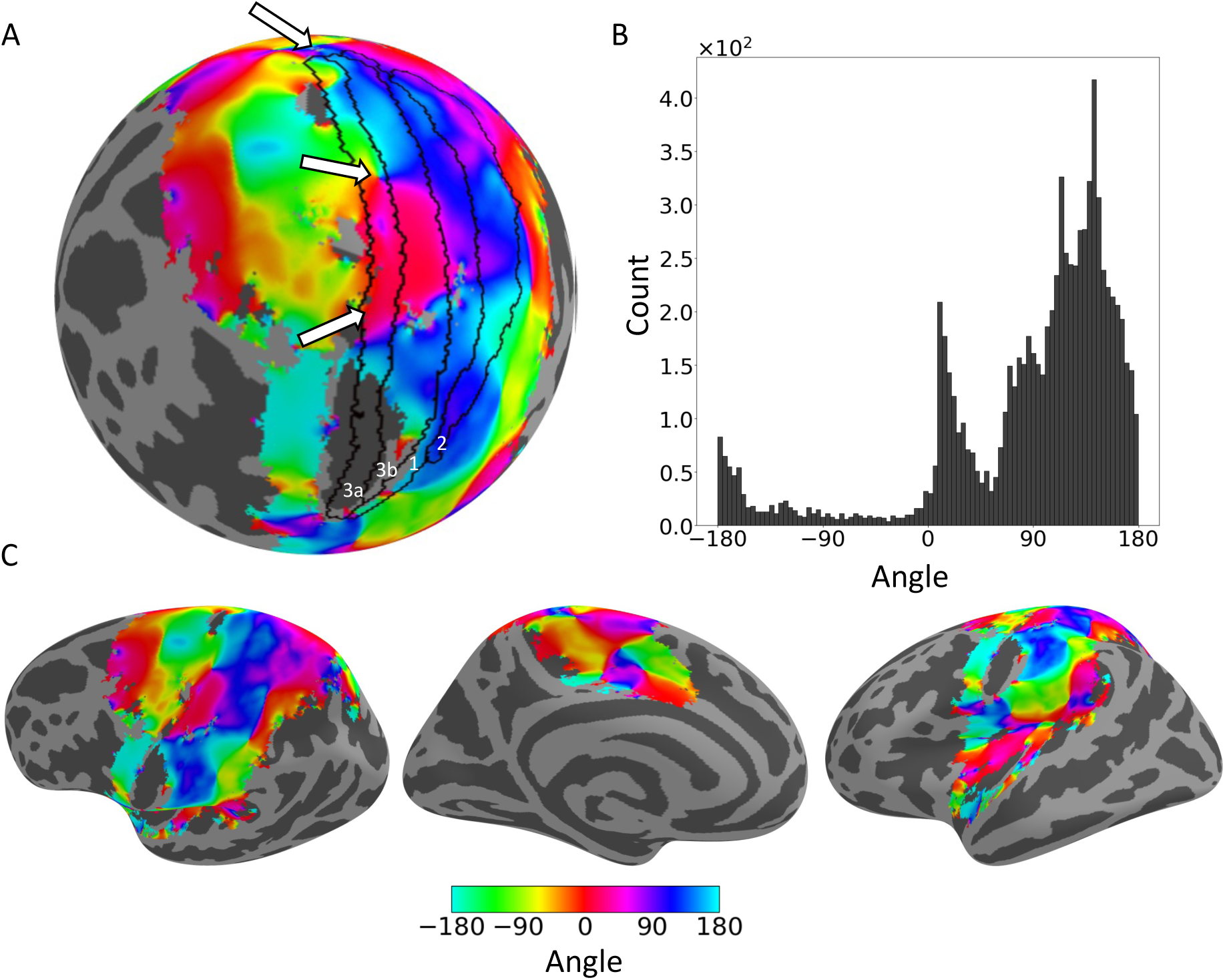
Geometric relationships of somatosensory body and hierarchy gradients. A. Color map of the angle between the local gradients of body and selectivity are displayed on a spherical representation of the left hemisphere cortical surface in a superior lateral view. Black lines denote the borders of Brodmann areas that comprise S1 (3a, 3b, 1, 2). White arrows mark the extrema of the two maps (see Fig. 3). Blue color that dominates the map within S1 borders indicates that the two gradients are approximately orthogonal. B. Histogram of the angles between the gradients within S1 with circular mean of *µ*_BS_ = 116° and circular variance of *V*_BS_ = 0.39. C. Color map of the angle between the local gradients of body and selectivity are displayed on an inflated surface in a lateral (left), medial (middle), and inferolateral (right) views. Note the variability across different cortical regions.

The blue-purple colors along S1 indicates that the body and selectivity gradients are, indeed, approximately orthogonal. However, the more quantitative histogram (Fig. 5B, S10B) reveals that the circular mean of the angle between the gradients of the body and selectivity maps is not exactly 90°. Rather, the circular mean is *µ*_BS_ = 116.4°. To quantify the width of the distribution, we used the circular variance. The circular variance is a measure of the width of a distribution of angles. It varies between 0 and 1. If all angles are the same (delta function distribution), *V* = 0; The circular variance of a uniform distribution over all angles is *V* = 1. If the distribution of angles is bimodal such that half of the angles are *θ* and half are *θ* + 180° then the circular variance is also maximal, *V* = 1. We found that the width of the distribution of angles between the body and selectivity gradient maps in S1 is *V*_BS_ = 0.38. For comparison, the circular mean and circular variance of the distribution of angles in S1 for the selectivity and laterality maps are *µ*_SL_ = 3.7° and *V*_*SL*_ = 0.07 (Figure S8). To understand the skewness of the distribution of angles, and the fact that circular mean *µ*_BS_ is substantially larger than 90°, we go back to the dipole metaphor, and note that S1 is not symmetric relative to the dipole. Rather, the maximum is at its border (medial extent) whereas the minimum is close to the center of S1 (in the medio-lateral direction). Thus, the distribution of angles contains regions lateral of the body minimum, in which the angles between the gradients are >90°, but not regions that are medial to the maximum, in which the angles are <90°.

We next asked whether the relationship between the gradients of the body and selectivity maps which we observed in S1 signifies an organizational principle that extends across the entire somatosensory-responsive cortex (Figs. 5C, S10C). We found substantial heterogeneity across the cortex. For example, in the anterior extent of the somatosensory cortex, in the frontal lobe, the gradients of the body and the hierarchy measures are parallel. In the precentral gyrus these gradients point at almost the opposite directions (−180°). In parietal regions, the angles between the gradients are distributed around 90°. Considering the entire distribution (Fig. S11), *µ*_BS_ = 8.37°. However, this circular mean is not very informative because the distribution is broad, with *V*_BS_ = 0.88. Taken together, the emerging picture is complex, and shows that the geometrical relationship between the body and hierarchy maps in S1 is not representative of that relationship throughout the entire somatosensory-responsive cortex.

## Discussion

In this study, we characterized the large-scale organization of the somatosensory cortex, focusing on the organization of body-part and hierarchy maps. We showed that on a large scale, the organization of both maps is radial, with a small number of maxima and minima that reign over a large number of cortical regions. Hierarchy is maximal in the anterior part of the postcentral gyrus. Two additional local maxima of hierarchy are found in the depth of the Sylvian fissure / posterior insula and in the superior medial wall. Full body representations originate from the lateral part of the pre- and post-central gyri superiorly, inferiorly and posteriorly. In S1, the body map changes in a direction that is approximately orthogonal to the change in hierarchy, allowing for the processing of the same body parts at multiple levels of hierarchy. However, this principle does not generalize to the entire somatosensory-responsive cortex. Rather, it is a pinhole observation of the large-scale radial geometry of the somatosensory representation in the cortex.

### Why radial geometry?

We can only speculate whether the radial geometry serves a particular computational or developmental function, or whether it is a byproduct of an evolutionary process. The function of radial hierarchy is, perhaps, easier to interpret. Maximal selectivity (the three maxima in our map) are found in and near the primary and secondary somatosensory cortices. From there, to all directions, selectivity decreases. These areas were shown to receive direct projections from the thalamus (Kaas 2012). Radial geometry of the body map is characterized by a lower-body ridge, surrounding a lips and hands basin. One can speculate that the fact that face and hands areas are localized, as opposed to the representation of legs that are sparsely represented along a ridge, allows the brain to process face and hands information more efficiently (shorter interconnections) than that of the legs. Indeed, the hands and face seem more important in somatosensation, as manifested, for example, by their higher discrimination ability (Catani 2017).

### Large-scale cortical organization in other sensory modalities

Large-scale radial topography is not unique to somatosensory processing and seems to characterize other sensory modalities. The visual field is retinotopically mapped in two dimensions-polar angle, and eccentricity. Visual representations in the cortex are organized into several clusters, groups of maps with parallel semicircular polar angle representations that share a common eccentricity representation (Wandell et al. 2005, 2007). The central foveal representation of each polar angle map is positioned in the center of the cluster and extends out to the periphery, spanning the radius radially, like a spokes on a wheel (Barton et al. 2012). The most obvious cluster in visual representation is the posterior cluster, centered on the occipital pole and includes V1, V2, V3 and hV4 representations (see Figure 9 in (Wandell et al. 2005)).

Similar radial topography was also suggested to characterize the auditory cortex (Barton et al. 2012). Auditory information is mapped in two dimensions-tonotopic, representing the spectral aspects of sound (i.e., tones) and periodotopic, representing the temporal aspects of sound (i.e., period or temporal envelope) (Barton et al. 2012; Brewer and Barton 2016). In each cluster, tonotopic representations runs from the center to the periphery of the cluster. Extending from the center of the cluster to its periphery are the periodotopic representations (see Figure 11 of (Brewer and Barton 2023)).

This type of radial topography, where a sensory variable is represented in radial bands from center to periphery of a cluster, may thus be a common organization to sensory systems (Brewer and Barton 2023).

### The geometrical relationship between body and hierarchy

In S1, the directions in which the body representation and selectivity change are nearly orthogonal. This orthogonality allows for a combinatorial coding in which every body part is represented at all levels of the hierarchy. However, if such orthogonality is functionally important, why doesn’t it generalize to the entire somatosensory cortex? Inspired by the functional organization of the visual cortex, we speculate that the answer to this question lies in the fact that aspects of information processing that go beyond simple body representation dominate the function of higher cortical regions (Longo et al. 2010; Tamè and Longo 2023). For example, higher-level somatosensory processing can be related to motor planning, visually guided movement, visceral signals and embodiment, and all these are also functionally mapped to the cortical surface. Because the cortex is a 2-dimensional sheet, it is impossible that all these functions will be orthogonal to the direction of hierarchical processing. We speculate that in higher cortical regions, these functions become orthogonal to hierarchy, at the expense of the body map.

The notion of functional specialization and different forms of hierarchy was suggested to characterize the visual system. The visual system consist parallel hierarchical sequences that are specialized for a particular functional task (Grill-Spector and Malach 2004). Early visual areas exhibit a high degree of retinotopy, but a low degree of specificity to a specialized functional task while higher-order areas show coarser retinotopy and a higher degree of specialization. Grill-Spector and Malach suggested that the large-scale organization of the visual cortex is governed by orthogonal axes of hierarchy and specialization.

### Limitations

There are several limitations to our approach in constructing body maps. First, we applied light touch using a brush. Somatosensory information is relayed to the brain using distinct “channels”, each corresponding to a different type of somatosensory information. Other types of somatosensory information are relayed via different fibers, and correspondingly, may form different body maps. The multiplicity of channels may indicate a possibility of a multiplicity of body maps, each corresponding to a different type of somatosensory information. For example, activity in Brodmann area 3a has been associated with proprioception (Krubitzer et al. 2004). This region selectively responded to touch in our experiments, but the magnitude of this response was relatively weak. Second, our stimulation was one-dimensional, along the long axis of the body. We did not study the representation in the orthogonal direction. Therefore, we only have partial information about the body parts representation. Finally, our study of face representation was limited. Due to technical limitations, our face stimulation was restricted to the lips. It is most likely that cortical regions that are more selective to other parts of the face, such as the forehead, nose, eyes and cheeks were registered as lips areas in our analysis.

With respect to hierarchy, hierarchy is a well-defined concept when information is processed serially, which (at best) only approximates information processing in the recurrent cortical network. Within this framework, its strongest prediction is that different regions within the hierarchy will be associated with a different response time to sensory stimulation. BOLD activation, however, does not provide the necessary temporal accuracy for such a measurement and our analysis was based on measures of selectivity (and laterality) that were used as proxies for the hierarchy level. Another limitation of hierarchy measurement is that it cannot be fully dissociated from the body map representation. For example, within S1, the hand and foot regions were found to be more selective in terms of receptive field size (Sur et al. 1980). Anatomical studies indicate that these regions are strongly contralateral, as ipsilateral information from the corpus callosum to these regions is almost absent (Iwamura 2000; Killackey et al. 1983). These can account for the existence of two maxima in and near area 3b, one in the anterior border of area 3b (in the hand area) and the other in the medial wall, near the medial border of area 3b (leg area). Notably, the inability to fully dissociate the hierarchy map from the body map exists even if one uses other measures of hierarchy. For example, consider response latency. Due to conductance speed, the delay from stimulation to response depends on the physical distance of the body part from the cortical region.

## Materials and methods

The collected data had previously been utilized for analyzing the spatial distribution of body parts (Saadon-Grosman et al. 2020b) and examining the hierarchical structure within the somatosensory responsive cortex (Saadon-Grosman et al. 2020a). Detailed experimental procedures are described there and in short below.

### Participants

All participants involved in this study provided written informed consent, and the study was approved by the ethical committee of the Hadassah Medical Center. 20 participants (age: 27.5 ± 3.33 year old, 9 females), all reported no history of neurological, psychiatric, or systemic health disorders.

### Somatosensory stimulation

A light-touch somatosensory stimulation was applied to the lips, dorsum part of the hand, forearm, upper arm, shoulder, trunk (lateral part), hip (lateral part), thigh (medial part), knee, shin (medial part), the dorsum part of the foot and the toes using a paintbrush (4 cm in width) by an experimenter, trained to maintain a constant pace and pressure hand (Saadon-Grosman et al. 2015, 2020a, 2020b; Tal et al. 2017). The stimulation was unilateral and continuous (without lifting the brush from the skin), except for one discontinuity between the lips and the hand. To control the timing of the body-part sequence, the experimenter wore fMRI-compatible headphones, delivering preprogrammed auditory cues. Stimulation duration was 15 s and the interval between stimulations was 12 s. Each scanning run included 7 repetitions of stimulation of one body side (right/left), followed by 7 repetitions of stimulation of the other body side (left/right). The order (right/left) was counter-balanced between participants. To control for time-order and directionality effect, BOLD activity of each participant was measured in two scanning runs that differed in the order of body-parts stimulations, from lips-to-toes and from toes-to-lips. Run duration was 423 s (282 time repetitions (TRs)), which included a 28.5 s of measurement before the onset of the first repetition and 4.5 s measurement after the last repetition, in addition to 12 s delay between the stimulation of the two body sides

### Functional MRI data acquisition

We used a Siemens Skyra 3T scanner, equipped with a 32-channel head coil, following a consistent imaging protocol. We utilized Blood Oxygen Level Dependent (BOLD) fMRI, employing a whole-brain gradient-echo echoplanar imaging sequence (GE-EPI). The specifics of this sequence included repetition time (TR)/time echo (TE) = 1500/27 ms, 90-degree flip angle, field of view (FOV) of 192×192 mm, and a 64×64 matrix yielding an in-plane resolution of 3×3 mm². The protocol involved acquiring 26 axial slices (slice thickness/gap = 4 mm/0.8 mm). Additionally, we acquired high-resolution T1-weighted anatomical images (1×1×1 mm) in the same orientation as the functional scans, with a TR/TE of 2300/2.98 ms, a 256×256 matrix, 160 axial slices, a 1-mm slice thickness, and an inversion time (TI) of 900 ms. For preprocessing, we utilized Freesurfer (’reconall’ for anatomical images (Fischl et al. 1999)) and FSL (’Feat’ for functional images (Jenkinson et al. 2012)), incorporating head motion correction, slice timing correction, and high-pass filtering with a 100-second cutoff. We also applied spatial smoothing with a Full Width Half Maximum (FWHM) of 4 mm. Each participant’s functional and anatomical data sets were co-registered (12 degrees of freedom, mcflirt, FSL), and projected to the fsaverage7 space (Freesurfer, 163,842 vertices per hemisphere). All subsequent analyses were executed using in-house custom scripts written in Python and Matlab.

### Identifying the somatosensory responsive cortex

To identify the somatosensory-responsive cortical regions, we employed a cross-correlation analysis method as described by Saadon-Grosman et al. (Saadon-Grosman et al. 2020a, 2020b). We conducted separate analyses for each side of the body. This involved dividing the vertex time course into segments of 137 TRs each (27 s per repetition for 7 repetitions, with 1.5 s per TR, resulting in 126 TRs plus 8 TRs before stimulation onset and 3 TRs post-stimulation). We used a boxcar function (3 s) convolved with a two-gamma hemodynamic response function (HRF) to create a predictor for our analysis. This predictor was then cross-correlated with the time course of each vertex to assess responses to various segments of the stimulation cycle. To reduce directionality effects, we used two stimulation directions, ‘start lips’ and ‘start toes’. The stimulation cycle was comprised of 10 TRs, resulting in 10 cross-correlations values. However, we excluded the first and last cross-correlations, as the first cross-correlation value may reflect a general response to the beginning of the stimulation block rather than the response to a specific body part. The last cross-correlation value was also excluded because later in the analysis, the order of the values of ‘start toes’ was reversed and averaged with the ‘start lips’ correlations (see below). A vertex was identified as significant for a participant if the maximal correlation value was r > 0.251, which corresponds to two-tailed t-test, α = 0.05, Bonferroni corrected for multiple correlations; 8 lag values × 2 directions, 135 degrees of freedom, P < 0.003. A vertex was considered significant for the group if it was significant for at least 13 of the participants (>2/3). Similar maps are obtained when applying random effect across participants (see Supplementary Fig. 1 in (Saadon-Grosman et al. 2020b)). Further refinement was conducted by eliminating clusters smaller than 3,000 vertices. The resulting final map, representing the somatosensory responsive cortex, was then used as a mask for subsequent analyses.

### Body map

To generate a body map, we averaged, for each vertex, the ‘start lips’ correlations with the ‘start toes’ correlations, time-reversed. The remaining 8 (out of 10) cross-correlations were assigned to a specific body part based on its stimulation time as follows: 1-lips, 2-distal upper limb, 3-proximal upper limb, 4-upper trunk, 5-lower trunk, 6-proximal lower limb, 7-mid-lower limb, 8-distal lower limb. These 8 cross-correlations were averaged over all 20 participants, resulting in 8 average correlation values which we denote by *r*_*i*_ where 1 ∈ {1, . . ., 8}. A naïve approach to identifying the body part associated with a vertex is to identify the highest correlation (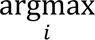_*i*_)). However, this approach would result in a discrete body map (8 discrete values). Because we were interested in computing local gradients, we developed an extrapolation method that provides a more refined and continuous body map. Specifically, rather than considering only the body part that corresponded to the highest correlation, we also considered the correlation values to the left and to the right of this value. For example, if the body part corresponding to the highest correlation was 5, we considered the correlation values with body parts 4, 5 and 6, *r*_4_, *r*_5_ and *r*_6_. We then fitted a parabola to these three values, and the corresponding body part was the location of the maximum of the fitted polynomial, which in our example would be a number between 4 and 6.

This approach cannot be used if the highest value is the first or last cross-correlation, because there is no adjacent value to the left or right, respectively. Consider, the case in which the highest value of the average correlation is 1 (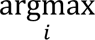(*r*_*i*_) = 1). The “true” body part will be between 0 and 2, but we do not have access to the correlation value associated with “0”. To overcome this limitation, we assumed that the distribution of body parts around 1, when the highest correlation is “1”, is comparable to the distribution of body parts around 2, when the highest correlation is “2”. Specifically, we computed, for each vertex in which argmax(*r*_*i*_) = 1) a measure of the steepness of the relative change from *r*_1_ to *r*_2_ : 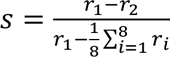. We expect that the larger the value of *s*, the larger the “true” value of the body part. We then computed the distribution of *s* over all the vertices where the highest correlation corresponded to body part “1”, and identified, for each vertex, its percentile rank. Next, we considered all vertices in which 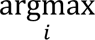(*r_i_*) = 2, and computed, for these vertices, the distribution of body parts (as computed by fitting a parabola to *r*_1_, *r*_2_ and *r*_3_). Finally, we posited that the body map value of a vertex for which 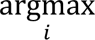(*r_i_*) = 1 is equal to the percentile of the vertex 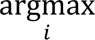(*r_i_*) = 2 that corresponds to the same percentile rank (its body map value), minus 1. The same procedure was repeated for all vertices in which 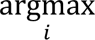(*r_i_*) = 8 in reverse, using the distribution of body maps values for all the vertices for which 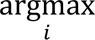(*r_i_*) = 7.

### Selectivity map

The method for computing the selectivity map follows our earlier work (refer to Saadon Grosman 2020 for detailed methodology) (Saadon-Grosman et al. 2020a): For each participant, in each stimulation direction and for each vertex, we normalized the time course (z-score) and computed event-related averaged response (ERA) across stimulation cycles (7 repetitions of 18 TRs each). We then fitted a Gaussian curve to the ERAs, *f*(*x*) ∝ 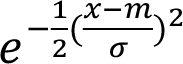, where *x* represents the ERA response time points (TRs), *σ* the standard deviation, and *m* the mean. We defined selectivity as s = √2⁄*σ* and averaged this measure over all participants and directions. Before averaging, we excluded from the analysis participants x direction in which the ERA was not well-fitted by a Gaussian (R2=residual sum of squares/total sum of squares; R2 < 0.6).

### Laterality map

The generation of the laterality map is described in Saadon Grosman et al., 2020 (Saadon-Grosman et al. 2020a). In short, we defined, for each participant, the laterality of a vertex as its preference for stimulation of either the contra- or ipsilateral body side. Denoted by *l*, the normalized difference between maximal responses to contra- and ipsilateral stimulation,

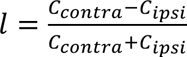, where *C* denotes the maximal correlation coefficient over the 8 body parts in the corresponding body side. Positive values of *l* indicate a preference for contralateral stimulation, whereas negative values of *l* indicate a preference for ipsilateral stimulation. Finally, we averaged the laterality values over all participants for whom the response in that vertex was statistically significant.

### Spatial Smoothing

Since measurements at the level of single vertices are noisy, computing local gradients requires spatial smoothing that would average out some of this noise. However, smoothing inevitably leads to information loss. Therefore, the magnitude of smoothing should consider the signal-to-noise ratio (SNR) at the level of individual vertices. The larger the SNR, the less smoothing is required. Considering the three maps (body, selectivity and laterality), we estimated the SNR for each map in the following way: the map value *x_i_* at each vertex 1 can be written as the sum of two terms, a signal term *µ_i_* and a noise term *ξ_i_*, *x_i_* = *µ_i_* + *ξ_i_*. In this framework, the SNR of the map is the ratio of the magnitude of a measure of the range of *µ*_*i*_ to the range of *ξ*_*i*_. Formally, we define 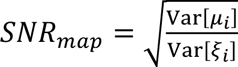. We have no direct access to *SNR*_*map*_ because we do not have an independent measure of *µ*_*i*_ and *ξ*_*i*_. However, we can estimate it in the following way: If *µ*_*i*_ and *ξ*_*i*_ are independent, then Var[*x*_*i*_] = Var[*µ*_*i*_] + Var[I_*i*_]. The total variance can be estimated by computing the variance of the map value over all the vertices. Assuming that the signal does not change much across neighboring vertices, Var[*ξ*_*i*_] can be estimated by calculating the variance over neighboring vertices Var[*ξ*_*i*_] ≈ Var[*x*_*i*_|*x*_*i*_ *are nearby*]. Therefore, to estimate a map’s SNR, we calculated, for each vertex the variance of it, and its nearest neighbors (7 vertices), and averaged this result over all vertices, E[Var[*x*_*i*_|*x*_*i*_ *are nearby*]]. The resultant SNR is estimated as: 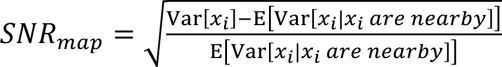.

We found that the SNR of the body, selectivity and laterality maps is *SNR*_body_ = 7.1, *SNR*_selectibity_ = 2.3 and *SNR*_laterality_ = 6.1, respectively. This result implies that the body map requires the least smoothing, whereas the selectivity map is the noisiest.

To smooth the maps, we recalculated the value of each vertex by averaging it with the values of its adjacent vertices (6 first-degree neighbors), repeating this process *n* times for each map. For a two-dimensional plane, this procedure corresponds to convolving the map with a 2-dimensional Gaussian filter, where the width is proportional to √*n*. In other words, the number of iterations *n* is effectively like averaging over *n* nearest neighbors, increasing the SNR by √*n*. We used *n* = 200, *n* = 800 and *n* = 400 for the body, selectivity, and laterality maps, respectively.

### Contour lines

To plot contour lines, we considered the distribution of map values. Vertices whose values were in quantiles 2.5% → 3.75% + 6.25% × *n* where *n* ∈ {0,1, . . ., 15} were colored in black.

### Spatial gradients

The challenge in computing the local gradient is that the cortical surface is convoluted and does not reside on a plane. Our approach was to consider, for each vertex, the plane that is orthogonal to the normal at that vertex. Specifically, each vertex in Freesurfer’s Fsaverage7 is characterized by its three-dimensional coordinates, and the vertices are linked by triangles such that each vertex is a node in six triangular faces. The normal at a vertex is defined as the average of the six normals of its six corresponding faces. For each vertex, we then projected all its third-degree neighboring vertices (36 in total) on that plane. Finally, we used linear regression to find the local gradient, defined as the vector that best predicts the 37 map values on the plane (center vertex and 36 neighbors) from their coordinates.

### Quantifying distributions

To quantify the distribution of angles between gradients, we used circular statistics.

Specifically, the circular mean is defined as 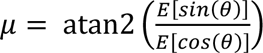 where *θ* denotes the angle between two gradients and the circular variance is defined as *V* = 1 −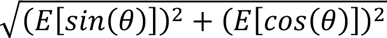.

### Identifying cortical regions

Cortical regions were defined using a multi-modal parcellation (Glasser et al. 2016). The regions were used to define S1 and its sub-regions (3a, 3b, 1, 2) and to orient in the different cortical surface representations (V1, A1, 4, 6, 7).

### Data and code availability

All data are available in the *Open neuro* repository (https://openneuro.org/datasets/ds003089/versions/1.0.1) and code in GitHub (https://github.com/tsahiasher/somatoGeometry)

## Acknowledgments

We thank Shahar Arzy for his support and Matthias Kaschube for fruitful discussions. We thank Simon Eickhoff for his critical insights regarding the representation in the secondary somatosensory cortex. We thank the ELSC MRI unit, Assaf Yohalashet, Lee Ashkenazi and Yuval Porat for their dedicated work.

