## Supporting information for "On the Geometry of Somatosensory Representations in the Cortex"

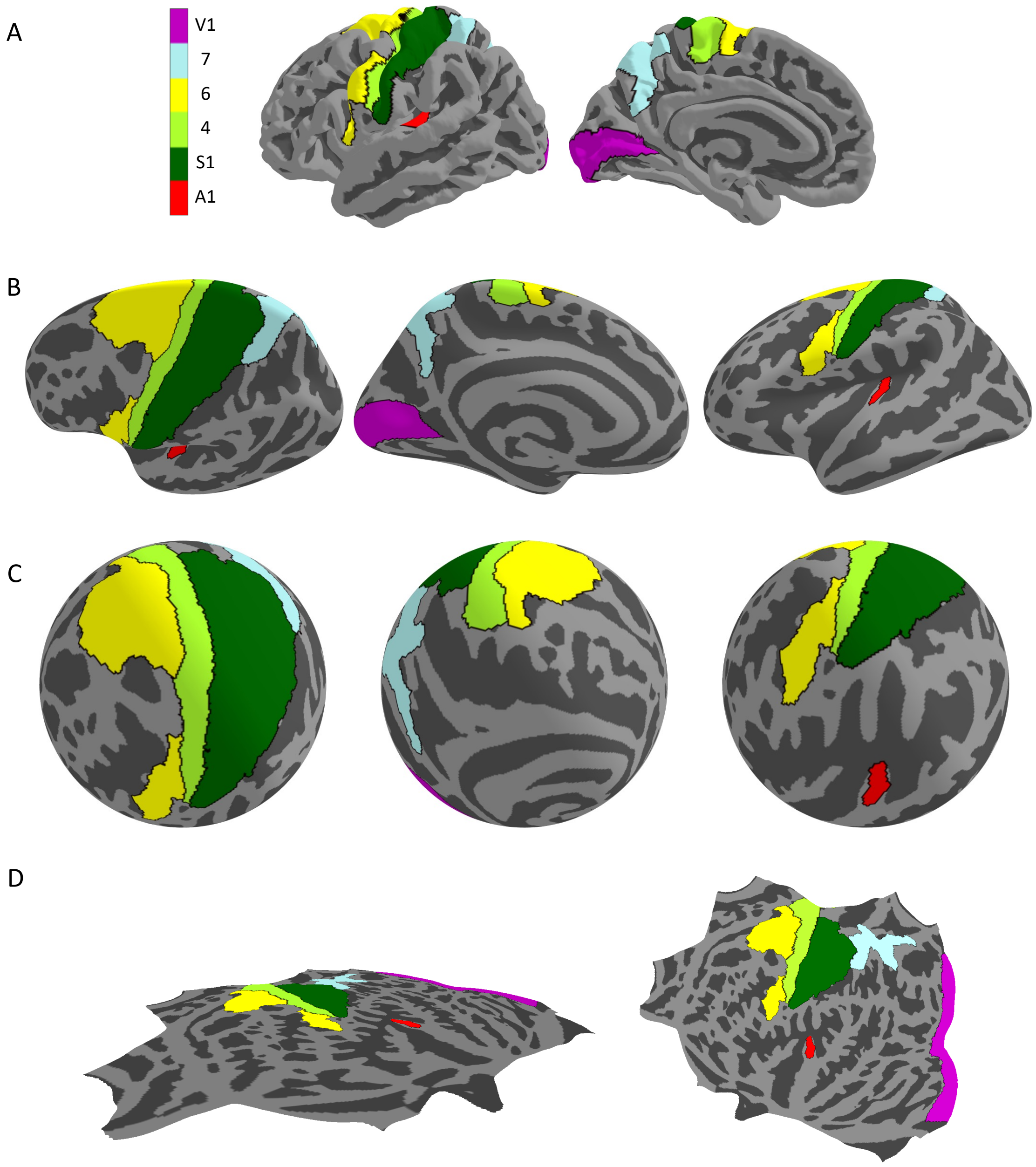


**Figure S1. Orientation in different surface representations of the left hemisphere**. To gain orientation within the cortical surface representations and their multiple views, established anatomical / functional areas are projected as follows: V1- primary visual cortex (pink), A1- primary auditory cortex (red), S1- primary somatosensory cortex (dark green), M1- primary motor cortex (light green), area 6- pre-motor cortex (yellow) and area 7- superior parietal (light blue). A. Pial surface in lateral (left) and medial (right) views. B. Inflated surface in lateral (left), medial (middle), and inferolateral (right) views. C. Spherical representation of the cortical surface in three views. D. Flat surface representation, showing a side (left) and a top (right) view. Areas were defined using a multi-modal parcellation by Glasser et al. (2016).

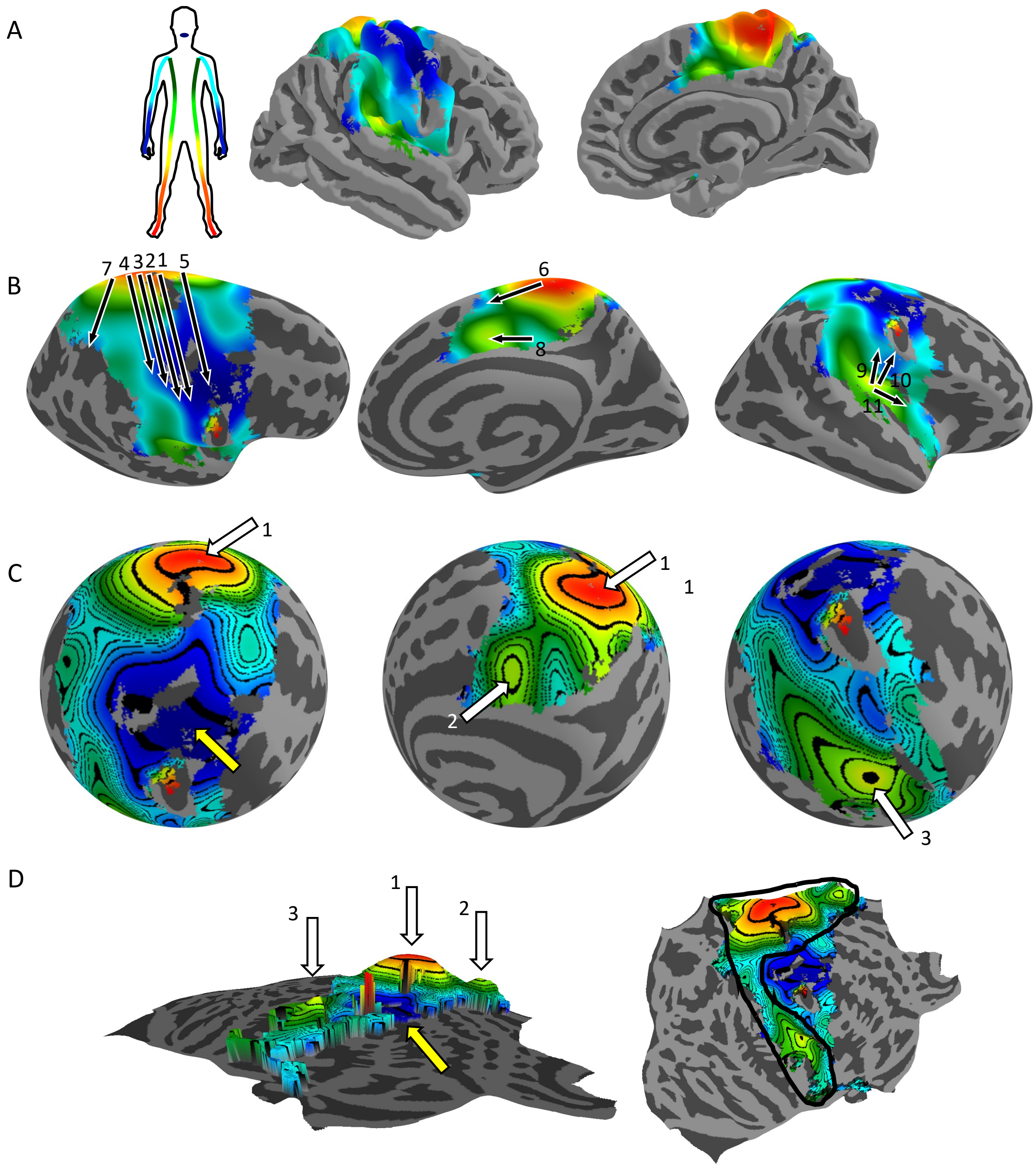


**Figure S2. Radial topography of somatosensory body representation- right hemisphere.** A group (N=20) body map corresponding to the stimulation of the contralateral body side, from lips (blue) to toes (red), is displayed on multiple representations of the right hemisphere cortical surface. A. Pial surface in lateral (left) and medial (right) views. B. Inflated surface in lateral (left), medial (middle), and inferolateral (right) views. Black arrows mark known body maps from previous literature (see text), note their consistency with the underlying map. C. Spherical representation of the cortical surface in three views. Contour lines represent vertices corresponding to the same body part. D. Flat surface representation, showing a side view where the body part value determines the height above the surface (left), and a top view of the same representation (right). The body map is characterized by a single minimum in the lateral part of the pre- and post-central gyri (lips, yellow arrow in C and D), surrounded by a semicircular ridge (black line in D) that spans the posterior extent of the entire somatosensory cortex on the medial and lateral surfaces. The ridge consists of three maxima (leg representation). The largest of these maxima lies at the midline, between the medial and lateral surfaces of the hemisphere, within the medial extent of the pre- and post-central gyri (arrow 1). A second, elongated peak is located deep within the Sylvian fissure/posterior insula (arrow 3). The third maximum is found in the cingulate gyrus on the opposite side of the ridge (arrow 2)

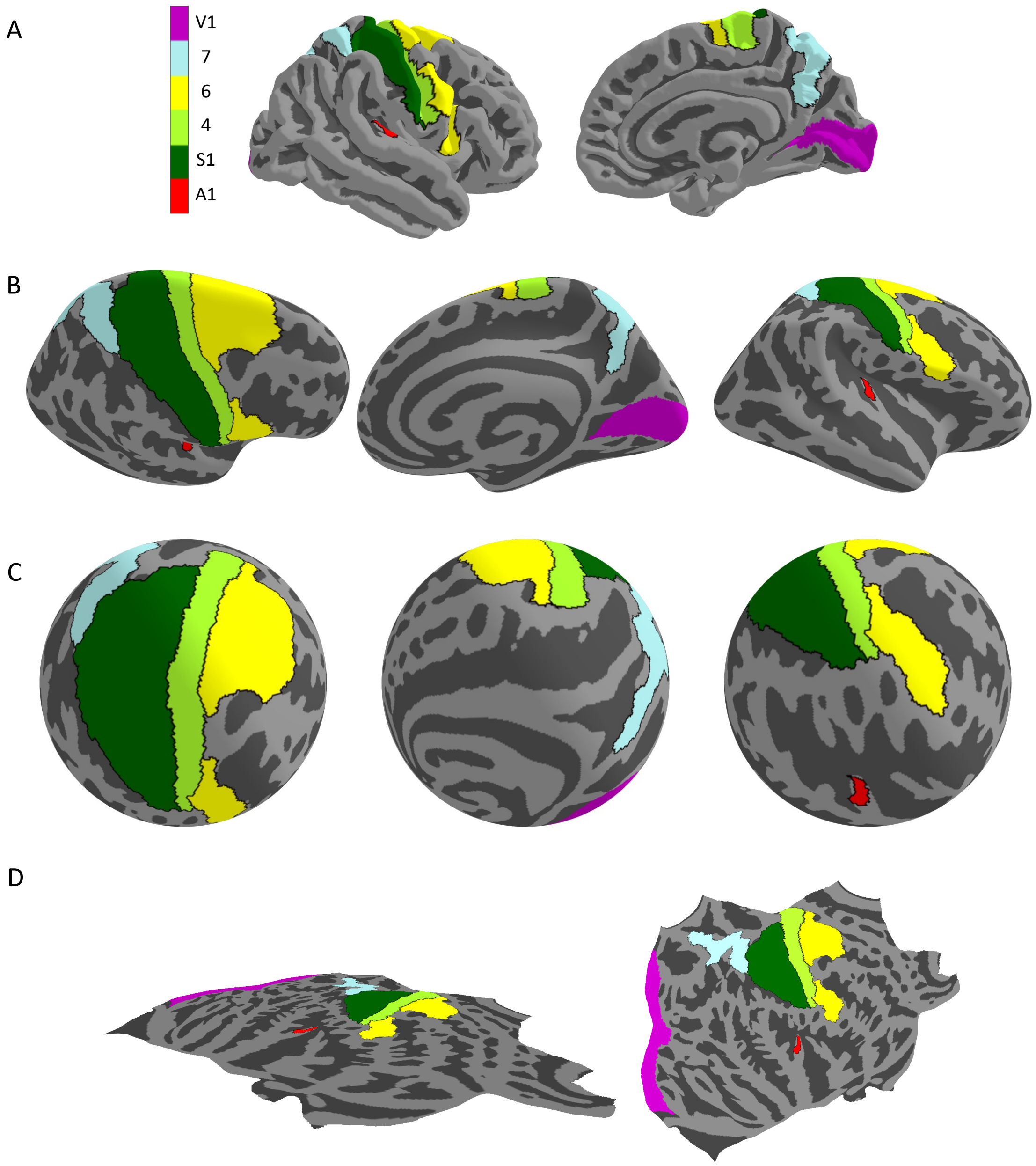


**Figure S3. Orientation in different surface representations of the right hemisphere**. To gain orientation within the cortical surface representations and their multiple views, established anatomical / functional areas are projected as follows: V1- primary visual cortex (pink), A1- primary auditory cortex (red), S1- primary somatosensory cortex (dark green), M1- primary motor cortex (light green), area 6- pre-motor cortex (yellow) and area 7- superior parietal (light blue). A. Pial surface in lateral (left) and medial (right) views. B. Inflated surface in lateral (left), medial (middle), and inferolateral (right) views. C. Spherical representation of the cortical surface in three views. D. Flat surface representation, showing a side (left) and a top (right) view. Areas were defined using a multi-modal parcellation by Glasser et al. (2016).

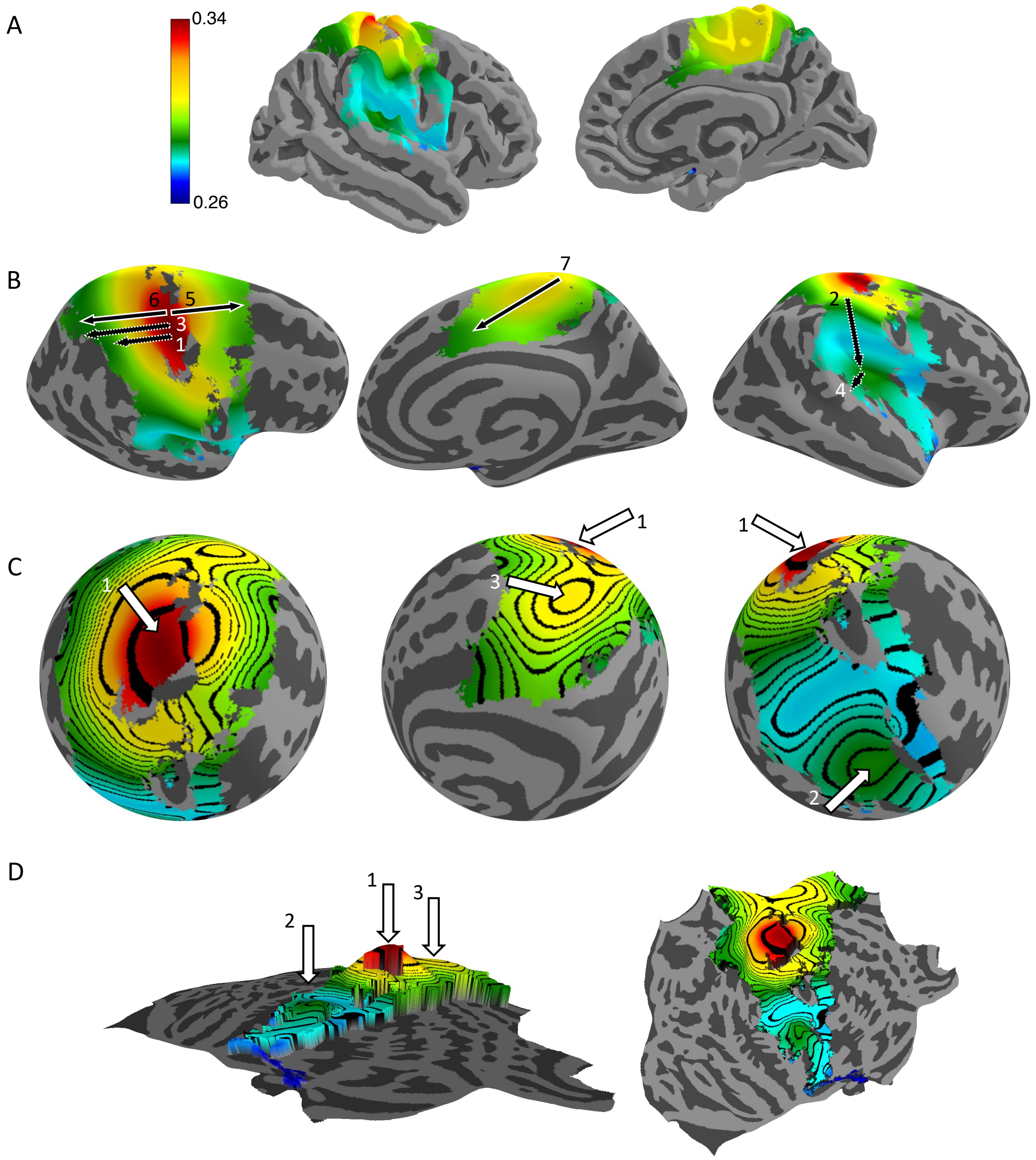


**Figure S4. Radial topography of somatosensory processing hierarchy- right hemisphere.** A group (N=20) selectivity map showing the specificity of the cortical response in each vertex to the preferred body part (high selectivity - red; low selectivity - blue) is displayed on various representations of the right hemisphere cortical surface (same as in Fig. 1). A. Pial surface in lateral (left) and medial (right) views. B. Inflated surface in lateral (left), medial (middle), and inferolateral (right) views. Black arrows indicate known hierarchies from previous literature (see text). Note the consistency of the arrows with the underlying selectivity map. C. Spherical representation of the cortical surface in three views. Contour lines represent vertices of equal selectivity. D. Flat surface representation, showing a side view where the selectivity value determines the height above the surface (left), and a top view of the same representation (right). The selectivity map is dominated by a maximum (a peak) in the anterior part of the postcentral gyrus (arrow 1). There are two additional local maxima of hierarchy: one in the depth of the Sylvian fissure/posterior insula (arrow 2), and one in the superior medial wall (arrow 3).

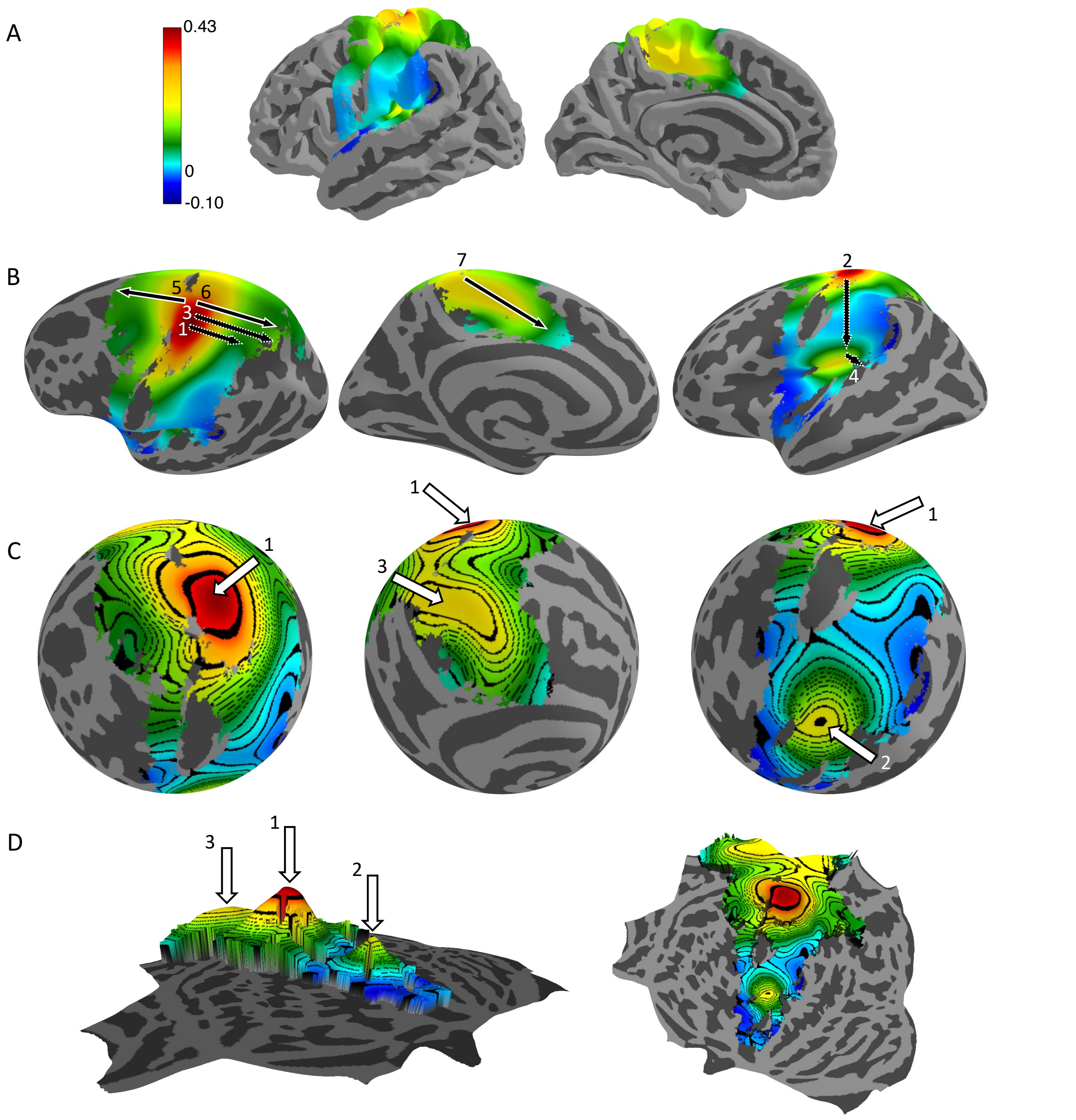


**Figure S5. Radial topography of somatosensory processing hierarchy- laterality, left hemisphere.** A group (N=20) laterality map showing the preference of each vertex to stimulation of the contra- or ipsi-lateral body-side (high laterality - red; low laterality - blue) is displayed on various representations of the left hemisphere cortical surface (same as in Fig. 2). A. Pial surface in lateral (left) and medial (right) views. B. Inflated surface in lateral (left), medial (middle), and inferolateral (right) views. Black arrows indicate known hierarchies from previous literature (see text). Note the consistency of the arrows with the underlying selectivity map. C. Spherical representation of the cortical surface in three views. Contour lines represent vertices of equal selectivity. D. Flat surface representation, showing a side view where the laterality value determines the height above the surface (left), and a top view of the same representation (right). The laterality map, as the selectivity map, is dominated by a maximum (a peak) in the anterior part of the postcentral gyrus (arrow 1). There are two additional local maxima of hierarchy: one in the depth of the Sylvian fissure/posterior insula (arrow 2), and one in the superior medial wall (arrow 3).

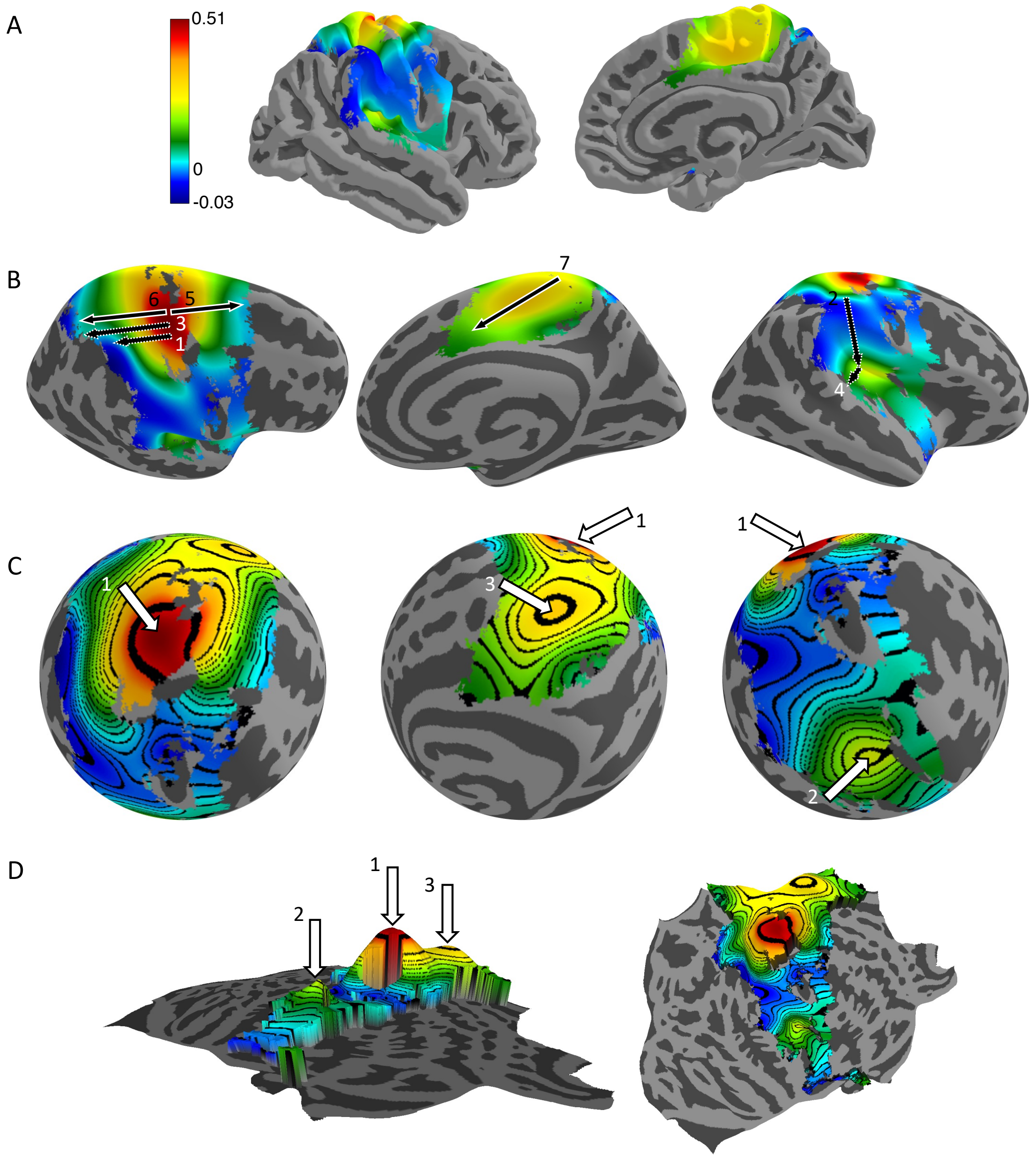


**Figure S6. Radial topography of somatosensory processing hierarchy- laterality, right hemisphere.** A group (N=20) laterality map showing the preference of each vertex to stimulation of the contra- or ipsi-lateral body-side (high laterality - red; low laterality - blue) is displayed on various representations of the right hemisphere cortical surface (same as in Fig. 2). A. Pial surface in lateral (left) and medial (right) views. B. Inflated surface in lateral (left), medial (middle), and inferolateral (right) views. Black arrows indicate known hierarchies from previous literature (see text). Note the consistency of the arrows with the underlying selectivity map. C. Spherical representation of the cortical surface in three views. Contour lines represent vertices of equal selectivity. D. Flat surface representation, showing a side view where the laterality value determines the height above the surface (left), and a top view of the same representation (right). The laterality map, as the selectivity map, is dominated by a maximum (a peak) in the anterior part of the postcentral gyrus (arrow 1). There are two additional local maxima of hierarchy: one in the depth of the Sylvian fissure/posterior insula (arrow 2), and one in the superior medial wall (arrow 3).

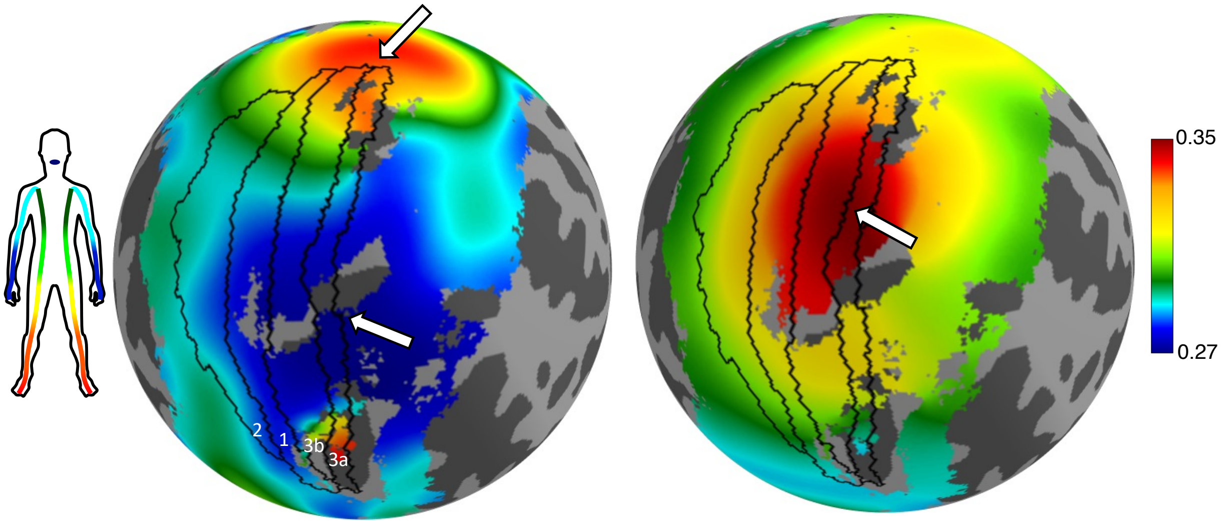


**Figure S7. Body representation and hierarchy in the primary somatosensory Cortex (S1)- right hemisphere.** Body map (left) and selectivity map (right) are displayed on a spherical representation of the right hemisphere cortical surface in a superior lateral view. Black lines denote the borders of Brodmann areas that comprise S1 (3a, 3b, 1, 2). Thick white arrows mark extrema. In the body map, a maximum (leg representation) is situated near the medial extent of S1, and a minimum (lips representation) is found in the lateral part of S1. In the selectivity map, a maximum is located approximately halfway between these body extrema.

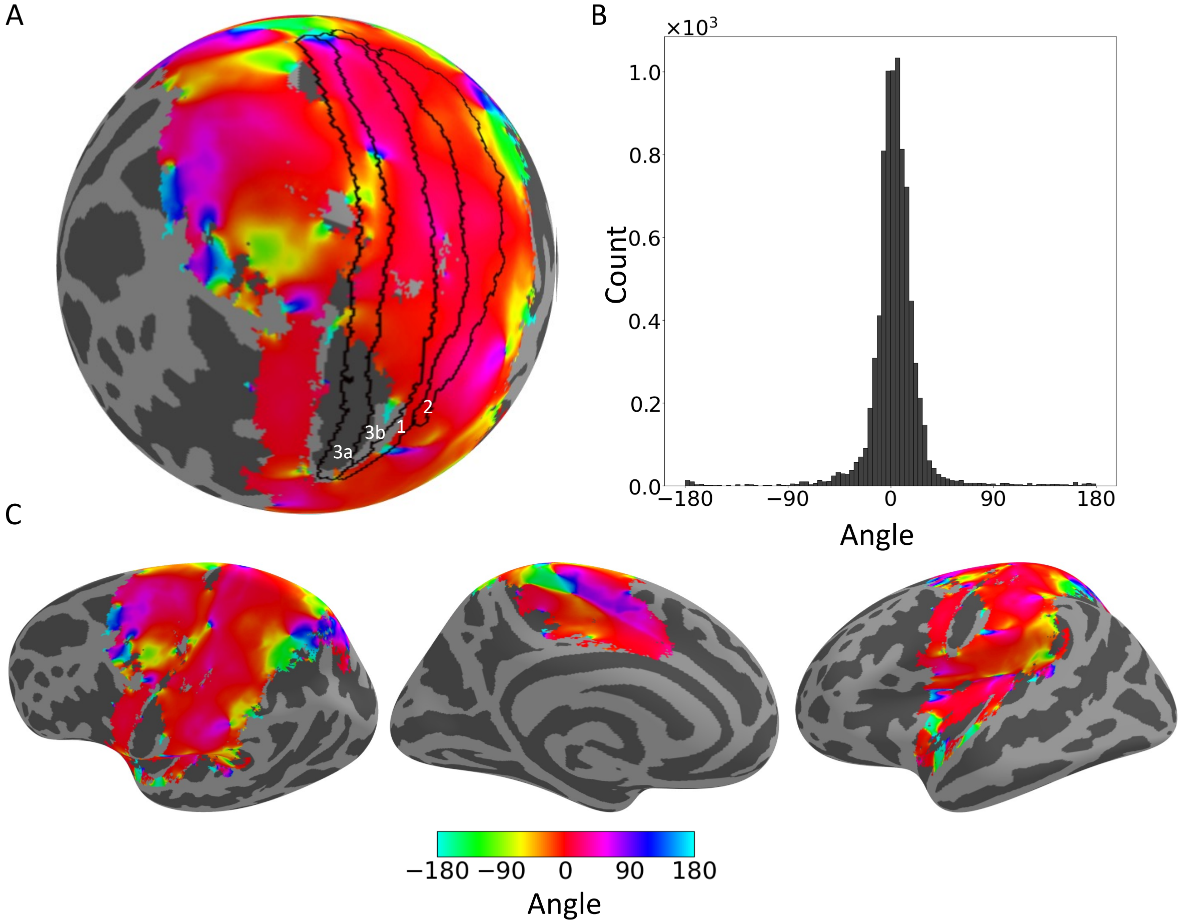


**Figure S8. Geometric relationships of somatosensory hierarchy gradients- left hemisphere. A**. Color map of the angle between the local gradients of selectivity and laterality are displayed on a spherical representation of the left hemisphere cortical surface in a superior lateral view. Black lines denote the borders of Brodmann areas that comprise S1 (3a, 3b, 1, 2). Red color that dominates the entire map indicates that the two gradients are approximately parallel. B. Histogram of the angles between the gradients within S1 with circular mean of $\mu_{BS} = 3.7°$ and circular variance of $V_{BS} = 0.07$. C. Color map of the angle between the local gradients of selectivity and laterality are displayed on an inflated surface in a lateral (left), medial (middle), and inferolateral (right) views.

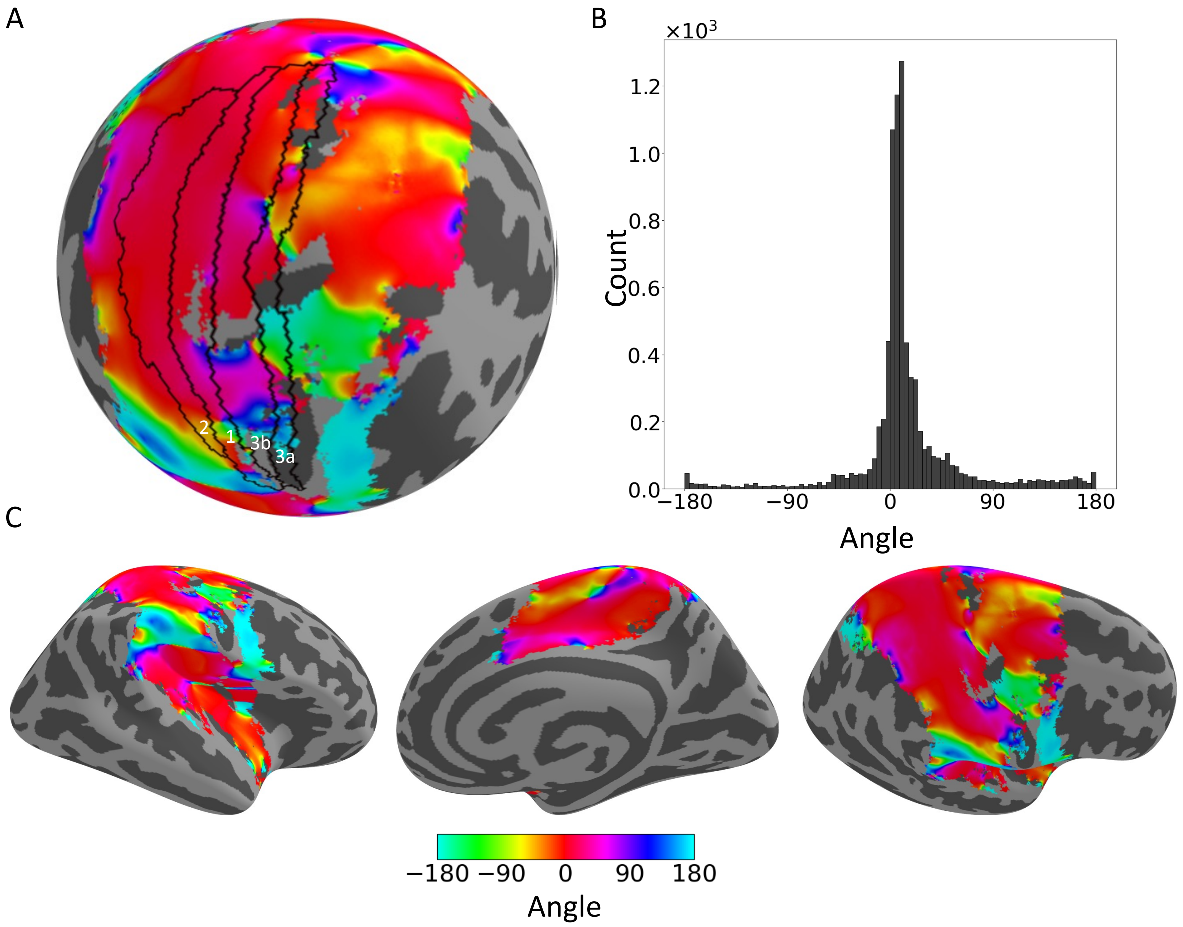


**Figure S9. Geometric relationships of somatosensory hierarchy gradients- right hemisphere. A**. Color map of the angle between the local gradients of selectivity and laterality are displayed on a spherical representation of the right hemisphere cortical surface in a superior lateral view. Black lines denote the borders of Brodmann areas that comprise S1 (3a, 3b, 1, 2). Red color that dominates the entire map indicates that the two gradients are approximately parallel. B. Histogram of the angles between the gradients within S1 with circular mean of $\mu_{BS} = 12.27°$ and circular variance of $V_{BS} = 0.24$. C. Color map of the angle between the local gradients of selectivity and laterality are displayed on an inflated surface in a lateral (left), medial (middle), and inferolateral (right) views.

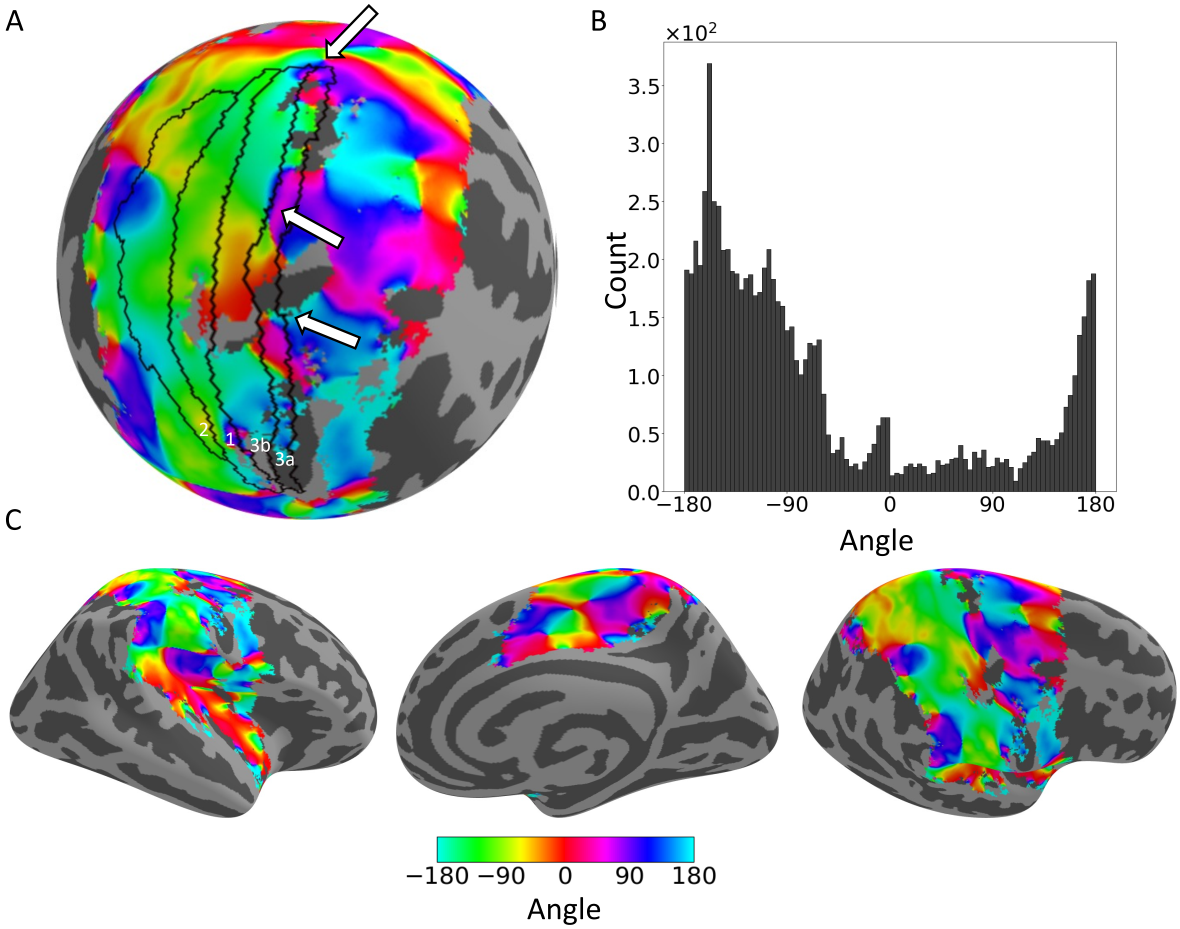


**Figure S10. Geometric relationships of somatosensory body and hierarchy gradients- right hemisphere. A**. Color map of the angle between the local gradients of body and selectivity are displayed on a spherical representation of the right hemisphere cortical surface in a superior lateral view. Black lines denote the borders of Brodmann areas that comprise S1 (3a, 3b, 1, 2). White arrows mark the extrema of the two maps (see Fig. 3). Green color that dominates the map within S1 borders indicates that the two gradients are approximately orthogonal. B. Histogram of the angles between the gradients within S1 with circular mean of $\mu_{BS} = -136.65°$ and circular variance of $V_{BS} = 0.46$. C. Color map of the angle between the local gradients of body and selectivity are displayed on an inflated surface in a lateral (left), medial (middle), and inferolateral (right) views. Note the variability across different cortical regions.

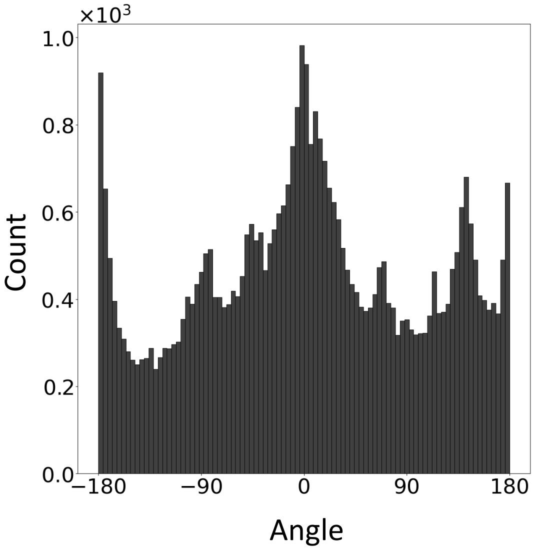


**Figure S11. Angles between somatosensory body and hierarchy gradients in the entire somatosensory responsive cortex.** Histogram of the angles between the gradients of body and selectivity in the entire somatosensory responsive cortex of the left hemisphere with circular mean of $\mu_{BS} = 8.37°$ and circular variance of $V_{BS} = 0.88$.
